# A new family of Type VI secretion system-delivered effector proteins displays ion-selective pore-forming activity

**DOI:** 10.1101/676247

**Authors:** Giuseppina Mariano, Katharina Trunk, David J. Williams, Laura Monlezun, Henrik Strahl, Samantha J. Pitt, Sarah J. Coulthurst

**Affiliations:** Division of Molecular Microbiology, School of Life Sciences, University of Dundee, Dow St, Dundee, DD1 5EH, UK; Centre for Bacterial Cell Biology, Newcastle University, Richardson Road, Newcastle-upon-Tyne, NE2 4AX, UK; School of Medicine, University of St Andrews, North Haugh, St Andrews KY16 9TF, UK

## Abstract

Type VI secretion systems (T6SSs) are nanomachines widely used by bacteria to compete with rivals. T6SSs deliver multiple toxic effector proteins directly into neighbouring cells and play key roles in shaping diverse polymicrobial communities. A number of families of T6SS-dependent anti-bacterial effectors have been characterised, however the mode of action of others remains unknown. Here we report that Ssp6, an anti-bacterial effector delivered by the *Serratia marcescens* T6SS, is an ion-selective pore-forming toxin. *In vivo*, Ssp6 inhibits growth by causing depolarisation of the inner membrane of intoxicated cells and also leads to increased outer membrane permeability, whilst reconstruction of Ssp6 activity *in vitro* demonstrated that it forms cation-selective pores. A survey of bacterial genomes revealed that Ssp6-like effectors are widespread in Enterobacteriaceae and often linked with T6SS genes. We conclude that Ssp6 represents a new family of T6SS-delivered anti-bacterial effectors, further diversifying the portfolio of weapons available for deployment during inter-bacterial conflict.

---

Bacteria have developed a variety of strategies to overcome their competitors and access limited resources, enabling them to survive and proliferate in multitude of polymicrobial environments. In some cases, these strategies involve actively killing or inhibiting the growth of rival bacteria. One mechanism widely used by Gram-negative bacteria for this kind of active competition is the Type VI secretion system (T6SS). The T6SS is a contact-dependent nanomachine which delivers toxic effector proteins directly into neighbouring cells. Bacteria most commonly use the T6SS to attack competitor bacteria, but this versatile weapon can also be used for manipulating host cells, killing fungal competitors or scavenging metals^1^. The T6SS is a mechanical puncturing device related to several contractile injection systems including bacteriophages^2^. According to the current model^1, 3–5^, contraction of an extended cytoplasmic sheath anchored in a trans-membrane basal complex propels a cell-puncturing structure, comprising a tube of Hcp hexamers tipped by a VgrG-PAAR spike, out of the secreting cell and towards a target cell. The Hcp-VgrG-PAAR structure is decorated by effector proteins which can interact covalently or non-covalently with one of these components. The rapid and powerful contraction events lead to the breach of a target cell by the expelled puncturing structure, followed by release of effectors inside the target cell.

A number of anti-bacterial effectors delivered by the T6SS have been described. These include large and diverse families of peptidoglycan hydrolases, phospholipases, nucleases and NAD(P)^+^-glycohydrolases^1, 6–11^. Singly or in combination, they provide effective killing or inhibition of targeted bacterial cells. In order to prevent self-intoxication or intoxication by genetically-identical neighbouring cells, T6SS-deploying bacteria possess a specific immunity protein for each anti-bacterial effector. Immunity proteins are encoded adjacent to their cognate effector, reside in the cellular compartment in which the effector exerts its action, and typically work by binding to the effector and occluding its active site^1^. Whilst many anti-bacterial T6SS effectors now have demonstrated or predictable functions, there remain many others whose function is unknown or not yet fully characterised. Intoxication by two unrelated effectors, Tse4 in *Pseudomonas aeruginosa* and VasX in *Vibrio cholerae*, led to loss of membrane potential and these effectors were proposed to be pore-forming toxins^12, 13^, however such a mechanism has not yet been demonstrated or characterised *in vitro* for a T6SS effector.

The opportunistic pathogen, *Serratia marcescens*, has a potent anti-bacterial T6SS, which secretes at least eight anti-bacterial effector proteins in addition to two anti-fungal effectors^14–16^. Several of these anti-bacterial effectors are not related to previously-characterised effectors and have mechanisms that cannot be readily predicted, therefore they likely represent novel anti-bacterial toxins. Here, we report the detailed characterisation of one of these new effectors, Ssp6. We reveal that Ssp6 acts by causing depolarisation of the target cell cytoplasmic membrane *in vivo* and provide a mechanistic explanation for this observation by demonstrating the ability of Ssp6 to form cation-selective pores *in vitro*. Homologues of Ssp6 can be found in many species of Enterobacteriaceae, hence Ssp6 defines a new family of T6SS-delivered, ion-selective pore-forming toxins.

## Results

### Ssp6 is a T6SS-delivered anti-bacterial effector protein and Sip6 is its cognate, membrane-located immunity protein

Ssp6 (SMDB11_4673) was identified as a small effector secreted by the T6SS of *S. marcescens* Db10 in previous studies using a mass spectrometry approach. However, its mode of action, which is not readily predictable from sequence-based or structural prediction methods, was not determined^14, 15^. Ssp6 is encoded outside the main T6SS gene cluster and is not linked with any T6SS genes (Fig. 1a). Using a strain of *S. marcescens* Db10 carrying Ssp6 fused with a C-terminal HA tag encoded at the normal chromosomal location (Ssp6-HA), we confirmed that Ssp6 is secreted in a T6SS-dependent manner, similar to the expelled component Hcp (Fig. 1b). No candidate immunity protein for Ssp6 is annotated in the published genome sequence of *S. marcescens* Db11 (a streptomycin-resistant derivative of Db10)^17^. We identified a 204 bp open reading frame (*SMDB11_4672A*) immediately downstream and overlapping by four nucleotides with *ssp6*. This genetic context strongly suggested that the encoded protein, named Sip6, represented the cognate immunity protein (Fig. 1a). To investigate the ability of Sip6 to inhibit Ssp6-mediated toxicity, we tested the susceptibility of a mutant lacking Sip6 to T6SS- and Ssp6-dependent inhibition by the wild type strain. The Δ*ssp6*Δ*sip6 ‘*target’ strain was indeed sensitive to T6SS-delivered Ssp6 activity, showing a loss in recovery when co-cultured with a wild type ‘attacker’ compared with attacker strains lacking an active T6SS (Δ*tssE*) or Ssp6 (Fig. 1c). The inability of the Δ*ssp6* mutant to cause intoxication could be complemented by expression of Ssp6 *in trans*, while expression of Sip6 restored the resistance of the Δ*ssp6*Δ*sip6* mutant against the wild type (Supplementary Fig. 1a). To confirm that Ssp6 and Sip6 are directly responsible for toxicity and immunity, respectively, Ssp6 with or without Sip6 was artificially expressed in *E. coli*. Ssp6 was either directed to the periplasm of *E. coli* through fusion with an N-terminal OmpA signal peptide (sp-Ssp6), or allowed to remain in the cytoplasm. Whilst Ssp6 was only mildly toxic when present in the cytoplasm, its presence in the periplasm caused pronounced inhibition of growth (Fig. 1d). This toxicity was alleviated upon co-expression of Sip6, thus confirming the identification of Sip6 as the cognate immunity protein of Ssp6.

**Figure 1.**
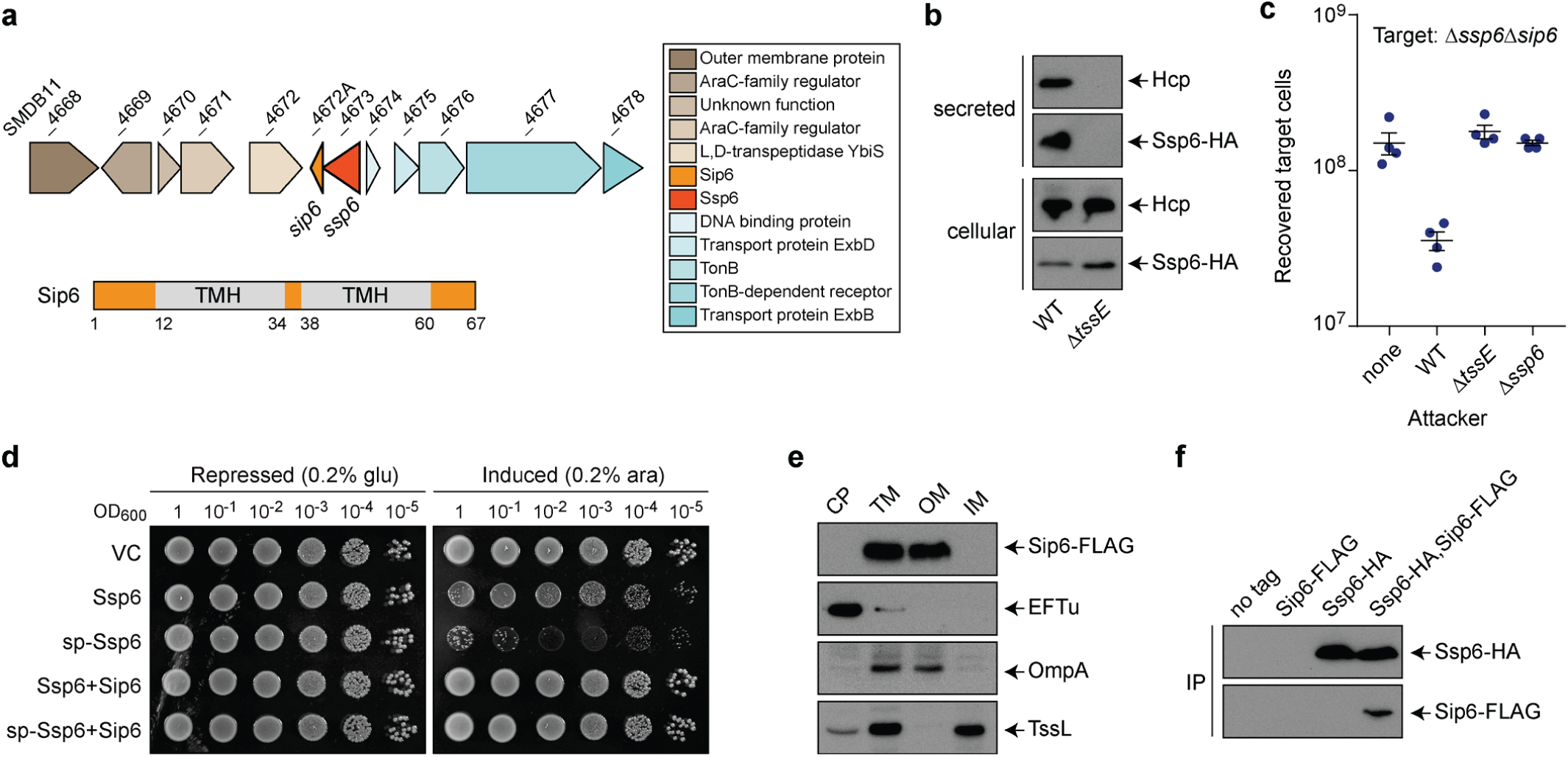
Ssp6 is a T6SS-delivered toxin and Sip6 is its cognate, membrane-associated immunity protein. (a) Schematic representation of the genomic context of the genes encoding Ssp6 and Sip6, with genomic identifiers (SMDB11_xxxx) provided above each gene and predicted protein functions in the box to the right. Below, the positions of the two transmembrane helices in Sip6, predicted using TMHMM v. 2.0, are indicated, where numbers refer to amino acids. (b) Immunoblot detection of Hcp1 and Ssp6-HA in cellular and secreted fractions of *S. marcescens* Db10 carrying the chromosomally-encoded Ssp6-HA fusion in either an otherwise wild type (WT) or T6SS-inactive (Δ*tssE*) background. (c) Number of recovered Δ*ssp6*Δ*sip6* target cells following co-culture with wild type (WT), Δ*tssE* or Δ*ssp6* mutant strains of *S. marcescens* Db10 as attackers. Individual data points are overlaid with the mean +/-SEM (n=4 biological replicates); none, target cells incubated with sterile media alone. (d) Growth of *E. coli* MG1655 carrying empty vector control (VC, pBAD18-Kn) or plasmids directing the expression of native Ssp6 (Ssp6) or Ssp6 fused with an N-terminal OmpA signal peptide (sp-Ssp6), each with or without Sip6, on LBA containing 0.2% D-glucose or 0.2% L-arabinose to repress or induce, respectively, gene expression. (e) Cells of *S. marcescens* Db10 carrying chromosomally-encoded Sip6-FLAG were subjected to subcellular fractionation and analysed by immunoblot detection of the FLAG epitope, EFTu (cytoplasmic control protein), TssL (inner membrane control protein) and OmpA (outer membrane control protein). CP, cytoplasm; TM, total membrane; OM, outer membrane; IM, inner membrane. (f) Co-immunoprecipitation of Ssp6-HA and Sip6-FLAG. Total cellular protein samples from wild type *S. marcescens* Db10 (no tagged proteins) and strains carrying chromosomally-encoded Ssp6-HA, Sip6-FLAG, or Ssp6-HA and Sip6-FLAG, were subjected to anti-HA immunoprecipitation and the resulting eluates were analysed by immunoblot.

In order to effectively prevent toxicity, T6SS immunity proteins are localised according to the cellular compartment in which the corresponding effector carries out its activity. Sip6 is predicted to contain two transmembrane helices (Fig. 1a), suggesting that Sip6 is localised in the membrane and that Ssp6 might intoxicate target cells by targeting their membranes. A strain of S. *marcescens* Db10 carrying a Sip6-FLAG fusion protein encoded at the normal chromosomal location was subjected to subcellular fractionation, which confirmed the presence of Sip6 in the membrane fraction (Fig. 1e). Interestingly, separation of the inner and outer membrane fractions revealed that Sip6-FLAG is localised in the outer membrane fraction. This was somewhat unexpected, since transmembrane helices are typically found in proteins that are localised in the inner membrane^18^, but is not unprecedented, since outer membrane proteins possessing α-helices rather than β-barrels have been described before^19^. Finally, to gain insight into how Sip6 neutralises Ssp6, a strain carrying both the chromosomal fusions Ssp6-HA and Sip6-FLAG was generated which exhibits full functionality for both Ssp6 toxicity and Sip6 immunity (Supplementary Figure 1c). This strain, together with control strains lacking either or both fusions, was used in a co-immunoprecipitation experiment. Sip6-FLAG was specifically co-precipitated with Ssp6-HA (Fig. 1f), demonstrating their interaction and suggesting that Sip6 acts directly on Ssp6 rather than by target protection or modification.

### Ssp6 intoxication induces stasis in target cells

To gain insight into the mode of action of Ssp6, we first aimed to determine whether it is a bacteriolytic effector, causing cell lysis, or a bacteriostatic effector, causing growth inhibition of target cells. We observed that artificially targeting Ssp6 to the periplasm of *E. coli* by inducing the expression of sp-Ssp6 resulted in cessation of growth but no drop in optical density, suggesting that Ssp6 is not bacteriolytic. When the inducer was removed, growth resumed and eventually reached the same optical density as control cultures lacking the toxin or co-expressing Sip6 (Fig. 2a). Next, we determined whether the action of Ssp6 in the most physiologically-relevant context, namely when delivered into target cells by the T6SS of a neighbouring cell, also results in a bacteriostatic effect. Cells of wild type, Δ*tssE* and Δ*ssp6* strains of *S. marcescens* Db10 were mixed with the Ssp6-susceptible target (Δ*ssp6*Δ*sip6*) on solid media and the growth of attacker and target cells was analysed over 3 h using time-lapse fluorescence microscopy. In these conditions, target cells in contact with wild type attacker cells generally failed to proliferate and divide, whilst target cells in contact with attackers unable to deliver Ssp6 proliferated indistinguishably from the attacking cells (Fig. 2b). The Ssp6-dependent impact on cell numbers was quantified by determining the fold increase in attacker and target cell populations between 0 h and 3 h in each condition. This population growth was noticeably reduced in target cells co-cultured with wild type attackers compared with growth of the attacking cells and of target cells co-cultured with attackers unable to deliver Ssp6 (Fig. 2c). It is important to note that 100% inhibition of target cell growth was not expected, since not every contact with an attacker cell or T6SS firing event would necessarily result in productive effector delivery. Given also that no lysis events were observed, these single cell microscopy data are consistent with a Ssp6 intoxication causing target cell stasis.

**Figure 2.**
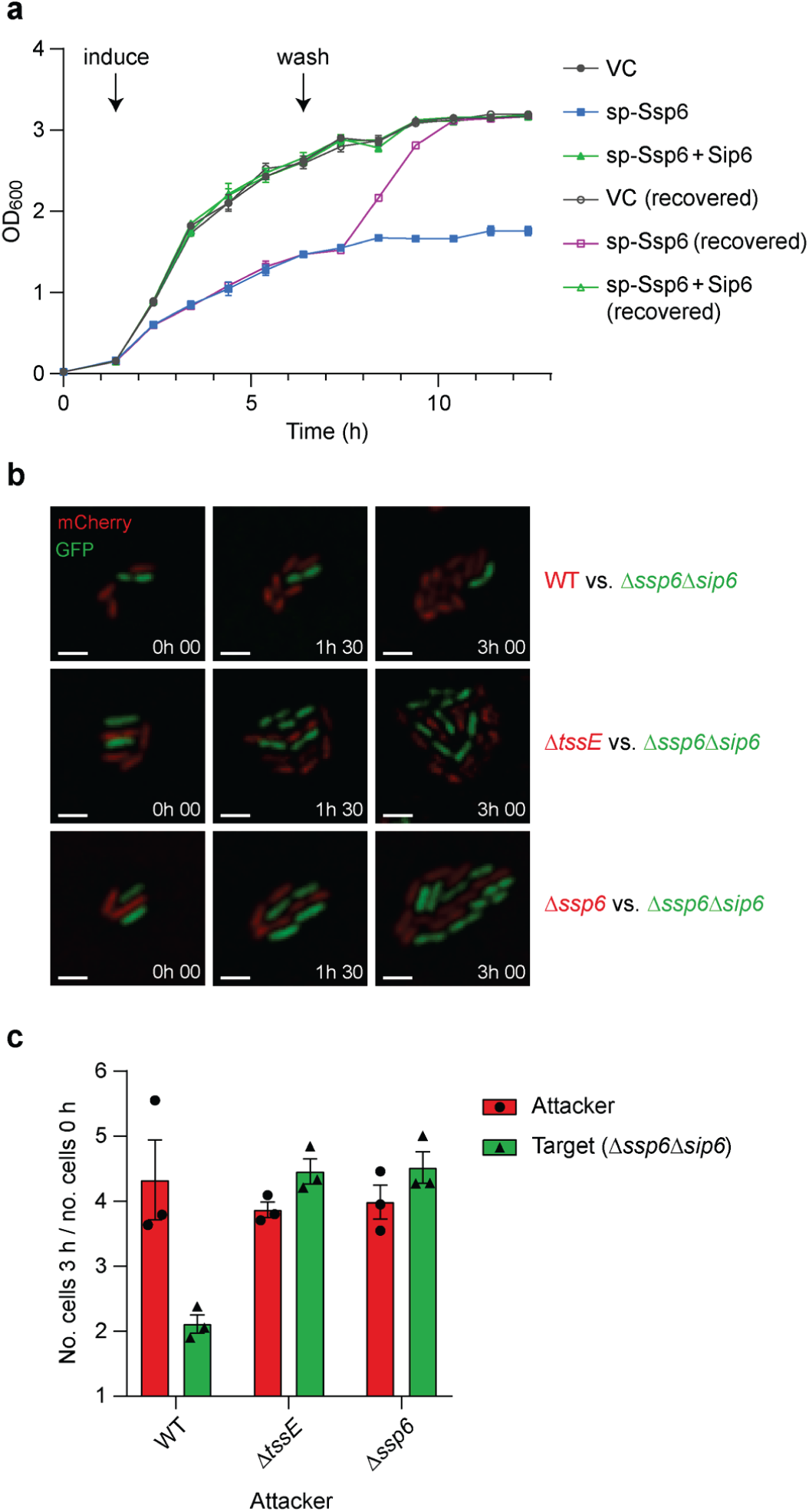
Intoxication by Ssp6 causes cessation of bacterial growth. (a) Growth in liquid LB media of *E. coli* MG1655 carrying empty vector control (VC, pBAD18-Kn) or plasmids directing the expression of Ssp6 fused with an N-terminal OmpA signal peptide (sp-Ssp6), either alone or with Sip6. To induce gene expression, 0.2% L-arabinose was added as indicated. To remove induction, the cells were washed and resuspended in fresh LB only (‘recovered’) at the ‘wash’ timepoint; control cells were resuspended in fresh LB + 0.2% L-arabinose. (b) Example images showing Ssp6-mediated growth inhibition as observed by time-lapse fluorescence microscopy. A Ssp6-susceptible target strain of *S. marcescens* Db10, Δ*ssp6*Δ*sip6* expressing cytoplasmic GFP (green), was co-cultured with wild type (WT) or mutant (Δ*ssp6* or Δ*tssE*) attacker strains expressing cytoplasmic mCherry (red) for 3 h. Scale bar 2 μm. (c) Quantification of time-lapse experiments. The total number of attacker cells and total number of target cells in at least ten microcolonies per experiment was counted at t = 0 h and 3 h and used to calculate fold increase in attacker and target cell numbers during the co-culture. Bars show mean +/-SEM, with individual data points superimposed (n=3 independent experiments).

### Ssp6-mediated toxicity causes depolarisation of target cells

The localisation of Sip6 suggested that the bacterial membrane might represent the target of Ssp6-mediated toxic activity. Since Ssp6 does not share any sequence or predicted structural similarity with phospholipase enzymes, we investigated whether Ssp6-mediated toxicity can affect the membrane potential and permeability of target cells. The impact of Ssp6 intoxication was analysed in the physiologically-relevant context, by co-culturing wild type, Δ*tssE* and Δ*ssp6* attacker strains of *S. marcescens* with the Ssp6-susceptible target, Δ*ssp6*Δ*sip6*, on solid media. Following co-culture, mixed populations of attacker and target cells were resuspended and stained with the voltage-sensitive dye DiBAC_4_(3). This negatively-charged dye is excluded from healthy, well-energised cells, resulting in low fluorescence. Upon membrane depolarisation, the dye can enter the cells and stain the cytoplasmic/inner membrane, causing an increase in green fluorescence^20^. Additionally, the samples were simultaneously stained with propidium iodide (PI), which cannot penetrate intact cells, but can enter cells with a damaged membrane, causing red fluorescence. Single cell analysis by flow cytometry revealed that just over 10% of the total population was depolarised when Δ*ssp6*Δ*sip6* was co-cultured with wild type. Of these cells, the majority showed disruption of membrane potential but no loss of membrane integrity (positive for DiBAC_4_(3) only, green fluorescence), whilst a small fraction showed both depolarisation and membrane permeabilisation (increased red and green fluorescence). In contrast, cells treated with polymyxin B, which causes formation of large, non-selective pores leading to cell permeabilisation and disruption of membrane potential^21^, never showed depolarisation (DiBAC_4_(3) fluorescence) without concomitant permeabilisation (PI staining). Depolarisation was specific to Ssp6 intoxication, since only background DiBAC_4_(3) and PI fluorescence was observed when Δ*ssp6*Δ*sip6* target cells were exposed to Δ*tssE* and Δ*ssp6* attackers (Fig. 3a, Supplementary Fig. 2). Whilst around 10% of the total population was depolarised, this total population contains both healthy attacker and Ssp6-intoxicated target cells. By considering both viable counts of recovered target cells (Fig. 1c) and counting of fluorescently labelled attacker and target cells (Supplementary Fig. 3) following co-cultures under the same conditions used in this experiment, we estimate that target cells represented approximately 20-35% of the total population following co-culture with the wild type. This suggests that Ssp6-mediated intoxication had caused detectable membrane depolarisation in around 1/3 of the target cells at the time of analysis. Finally, we confirmed that a similar pattern of membrane depolarisation was observed when Ssp6 was expressed in *E. coli*. Upon expression of sp-Ssp6, around half of the cells were depolarised, with the majority retaining gross membrane integrity (Fig. 3b). As expected, when Sip6 was co-expressed with sp-Ssp6, the number of DiBAC_4_(3) and PI positive cells was indistinguishable from that in control cells carrying the empty vector. Overall these results indicate that Ssp6 can disrupt the membrane potential of target cells in a mechanism that does not involve the formation of large unspecific pores or gross loss of inner membrane bilayer integrity.

**Figure 3.**
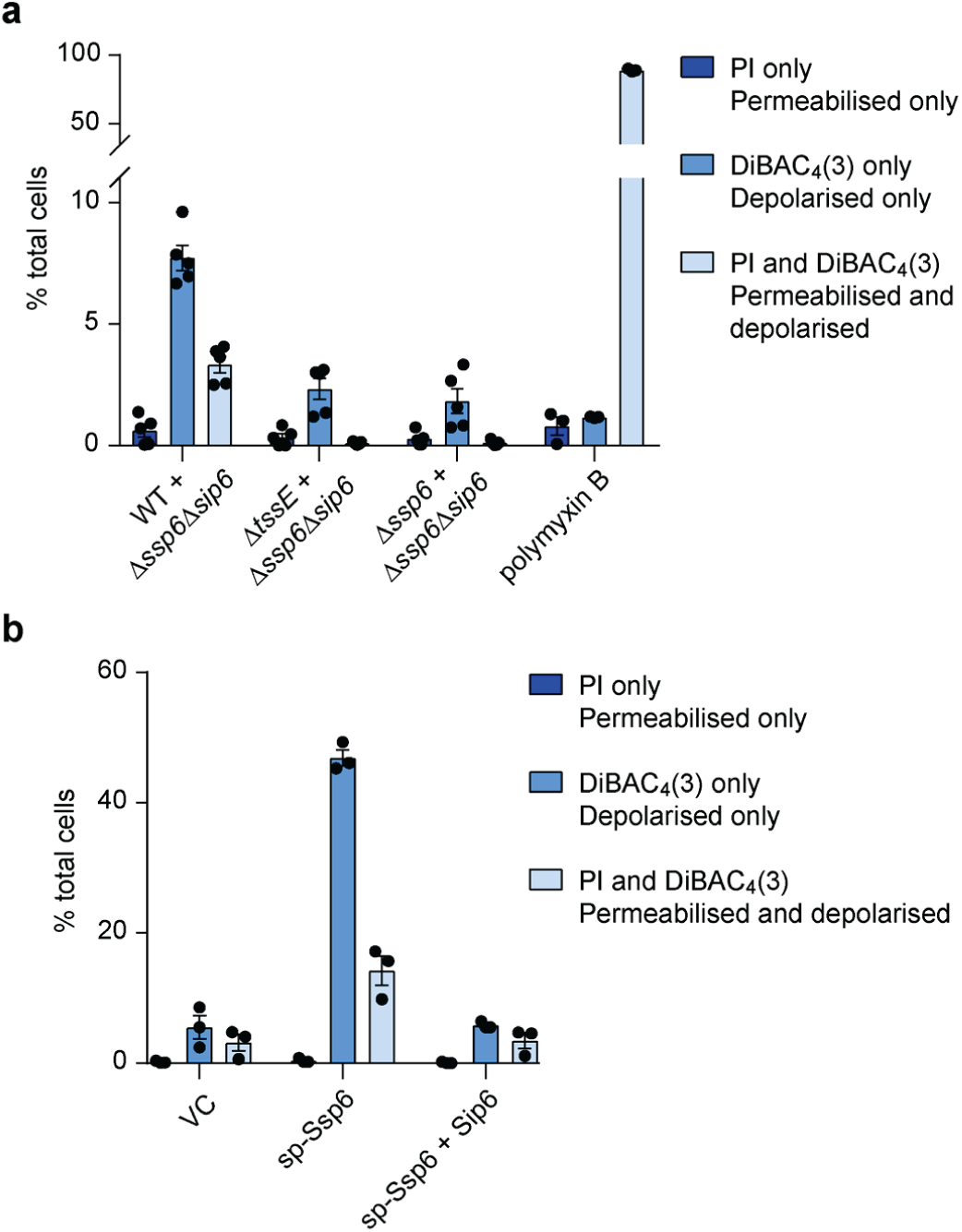
Ssp6 intoxication via T6SS delivery or heterologous expression causes loss of membrane potential. (a) The Ssp6-susceptible mutant of *S. marcescens* Db10, Δ*ssp6*Δ*sip6*, was co-cultured with the wild type (WT), Δ*tssE* mutant or Δ*ssp6* mutant and then membrane potential and membrane permeability of the mixed population was determined. Cells were stained with DiBAC_4_(3) and propidium iodide (PI) and analysed by flow cytometry, allowing different populations to be detected: depolarised (increased green fluorescence from DiBAC_4_(3)), permeabilised (red fluorescence from PI), depolarised and permeabilised cells (green fluorescence and red fluorescence), and healthy cells (below threshold fluorescence). The percentage of cells in the total mixed co-culture population identified as being permeabilised only, depolarised only, or simultaneously depolarised and permeabilised is shown on the Y-axis. Bars show mean +/-SEM, with individual data points superimposed (n=5 independent experiments, except for the polymyxin B control, where n=3). (b) Membrane potential and permeability of cells of *E. coli* MG1655 carrying empty vector control (VC, pBAD18-Kn) or plasmids directing the expression of Ssp6 fused with an N-terminal OmpA signal peptide (sp-Ssp6), either alone or with Sip6, was determined as in part a. Bars show mean +/-SEM, with individual data points superimposed (n = 3 independent experiments).

### Ssp6 forms ion-selective membrane pores *in vitro*

Our observations *in vivo*, that Ssp6 intoxication can cause depolarisation of target cells without increasing membrane permeability for larger compounds such as PI, suggested that Ssp6 could act though the formation of an ion-selective pore, leading to ion leakage and disruption of membrane potential. To examine potential pore-forming properties of Ssp6, the protein was purified as a fusion with maltose binding protein (MBP) and found to form higher order oligomers (Supplementary Fig. 4a). The MBP-Ssp6 fusion protein retained toxic activity upon expression in *E. coli*, causing growth inhibition which was similar to Ssp6 alone and reduced by co-expression of Sip6 (Supplementary Fig. 4b).

To test the ability of Ssp6 to form pores, the purified MBP-Ssp6 protein was incorporated into artificial membranes, under voltage-clamp conditions in non-symmetrical conditions (210 mM KCl in the *trans* chamber and 510 mM KCl in the *cis* chamber). Incorporation of MBP-Ssp6 generated a current, thus revealing that Ssp6 could indeed form ion-conducting channels. We observed that the Ssp6-mediated pore could exist in different opening states (Supplementary Fig. 5ab) and that openings and closings of the pore were too rapid to accurately measure gating properties. For this reason, we used noise analysis to determine the ion selectivity. To investigate whether Ssp6 is permeable to cations or anions, we tested if its reversal potential (calculated according to the Nernst equation) of the current/voltage (*I*/*V*) relationship in non-symmetrical conditions was shifted towards the equilibrium potential of K^+^ or Cl^−^. In these conditions, the reversal potential was −26.5 ± 4.4 mV, a value that is very close to the predicted equilibrium potential of potassium (−22.8 mV), indicating that Ssp6 can form a pore that shows a strong preference for cations, although a small contribution of Cl^−^ cannot be excluded (Fig. 4a). As controls, we used the purification buffer alone and MBP alone and applied a holding potential of +50 mV. In these conditions, no currents were observed, confirming that pore formation can be attributed to Ssp6 (Supplementary Fig. 5cd).

**Figure 4.**
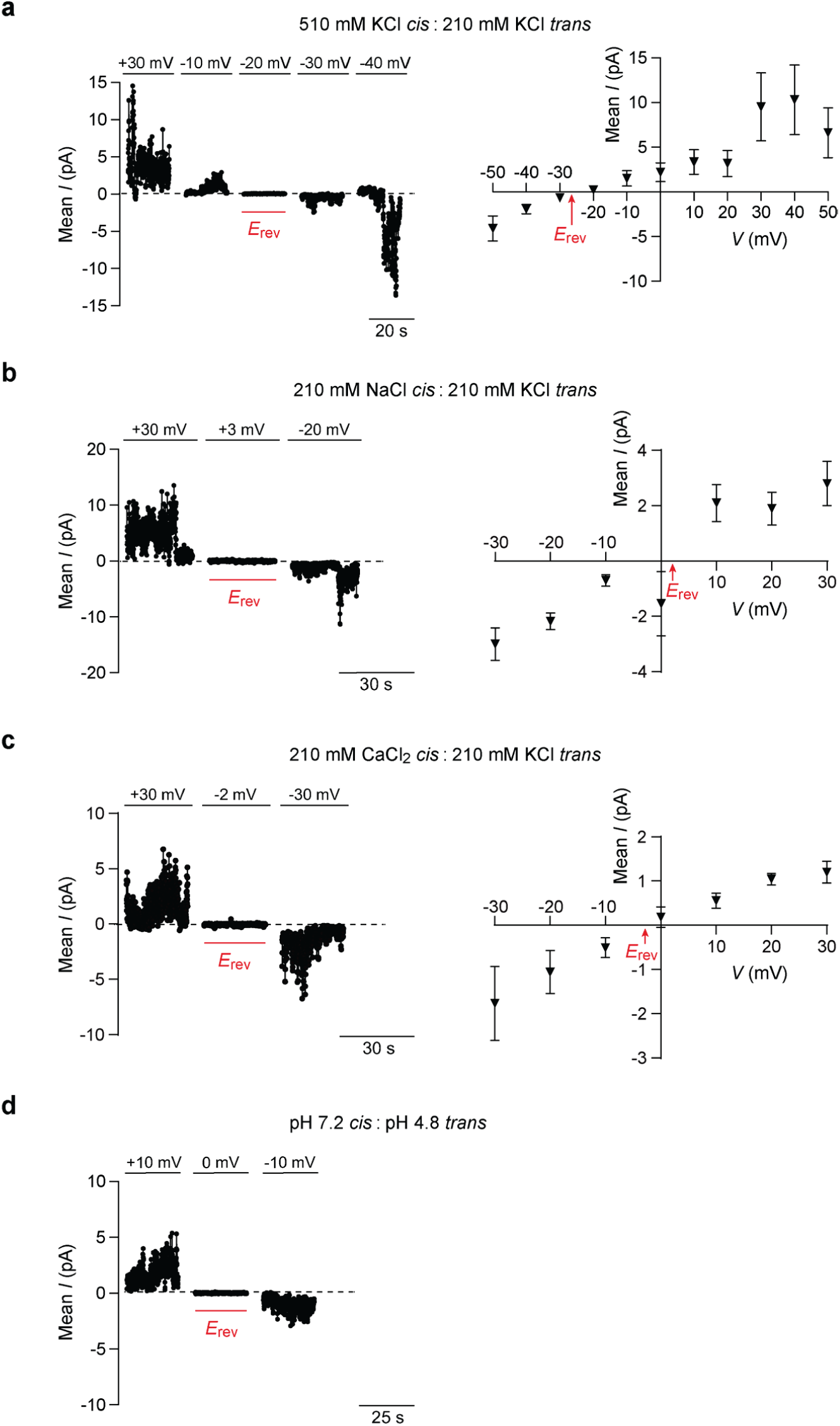
Ssp6 can form cation-selective pores *in vitro*. Multiple pores formed by MBP-Ssp6 were incorporated into the lipid bilayer under voltage clamp conditions and the mean current was measured. (a-c) Noise analysis of current fluctuations at the indicated holding potentials (left panels) and *I*/*V* relationship (right panels) for MBP-Ssp6 under conditions of: (a) a KCl gradient (510 mM *cis* chamber:210 mM *trans* chamber); (b) Na^+^ in the *cis* chamber (210 mM NaCl) and K^+^ in the *trans* chamber (210 mM KCl); and (c) Ca^2+^ in the *cis* chamber (210 mM CaCl_2_) and K^+^ in the *trans* chamber (210 mM KCl). (d) Noise analysis of current fluctuations at −10 mV, 0 mV and +10 mV under conditions of symmetrical 210 mM K^+^ acetate with the *cis* chamber at pH 7.2 and the *trans* chamber at pH 4.8. (a-d) The dotted line in each current trace shows the zero current level (left panels); points for *I*/*V* relationships (right panels) show mean +/-SEM (n=3 independent experiments, except part a where n=4); reversal potential (*E*_rev_) is indicated in red.

Given that Ssp6 displays a strong preference for cations, we next examined the relative permeability of K^+^, Na^+^ and Ca^2+^ to establish if Ssp6 has higher selectivity for monovalent or divalent cations. The relative K^+^/Na^+^ permeability ratio (*P*K^+^/*P*Na^+^) was assessed under conditions in which Na^+^ was the permeant ion in the *cis* chamber and K^+^ was the permeant ion in the *trans* chamber. In these conditions, the reversal potential was 2.07 ± 0.77 mV, corresponding to *P*K^+^/*P*Na^+^ of 0.93 ± 0.03, indicating that under bi-ionic conditions, Ssp6 has a similar selectivity for Na^+^ and K^+^ (Fig. 4b). To examine the relative K^+^/Ca^2+^ permeability, we used conditions in which K^+^ was the only permeant ion in the *trans* chamber and Ca^2+^ was the only permeant ion in the *cis* chamber. In this case, the reversal potential was −2.83 ± 2.08 mV and the relative K^+^/Ca^2+^ permeability ratio (*P*K^+^ /*P*Ca^2+^) was 8.80 ± 1.47, highlighting that the pore formed by Ssp6 is more selective for monovalent cations than divalent cations (Fig. 4c).

Finally, given that extrusion of protons (H^+^) out of the bacterial cytoplasm into the periplasmic space contributes to maintenance of a negative membrane potential, we tested whether the Ssp6 pore is permeable to H^+^. In this experiment, 210 mM potassium acetate pH 4.8 was present in the *trans* chamber and 210 mM potassium acetate pH 7.2 in the *cis* chamber. When the holding potential was 0 mV, no current was observed (Fig. 4d). In these conditions, the only permeant ions would be H^+^ and the driving force for its movement through the pore would be its chemical gradient. Given that we observed no current at 0 mV, our results indicate that Ssp6 pore is not permeant to H^+^.

### Ssp6 intoxication impairs the integrity of the outer membrane

In the previous sections we showed that Ssp6-mediated intoxication causes depolarisation of the inner membrane of target cells. However, we also observed that its cognate immunity protein, Sip6, is localised in the outer membrane. Therefore we investigated whether Ssp6 can also damage the outer membrane of target cells. First, we used the membrane-specific stain FM4-64, a lipophilic dye that can stain the outer membrane of Gram-negative bacteria^22, 23^. When cells of *E. coli* carried an empty vector, or sp-Ssp6 together with Sip6, the typical evenly-distributed fluorescence of FM4-64 outlining the cells was observed. However when sp-Ssp6 was expressed in *E. coli*, the red signal was not uniform in its distribution, showing a tendency to accumulate in ‘spots’, often at the cell poles (Fig. 5a). This experiment could not be performed using co-cultures of *S. marcescens* Db10, since this organism does not stain well with FM4-64 for reasons that are not known. Aberrant Ssp6-induced FM4-64 staining in *E. coli* cells suggested that Ssp6-mediated intoxication might alter the lipid organisation and thus, potentially, the integrity of the outer membrane. The fluorescent probe 1-*N*-phenylnapthylamine (NPN) was used to determine whether expression of sp-Ssp6 in *E. coli* increases the permeability of the outer membrane. NPN is unable to cross the outer membrane and displays weak fluorescence in aqueous solution. However, if the permeability of the outer membrane is increased, NPN can bind strongly to phospholipids, increasing its fluorescence^24, 25^. Expression of sp-Ssp6 in *E. coli* caused a large increase in NPN uptake, which was similar to the positive control EDTA but not observed in cells carrying the empty vector or co-expressing Sip6. Therefore Ssp6-mediated intoxication can cause an increase in the permeability of the outer membrane of target cells, which may be associated with the observed microscopic changes in FM4-64 staining.

**Figure 5.**
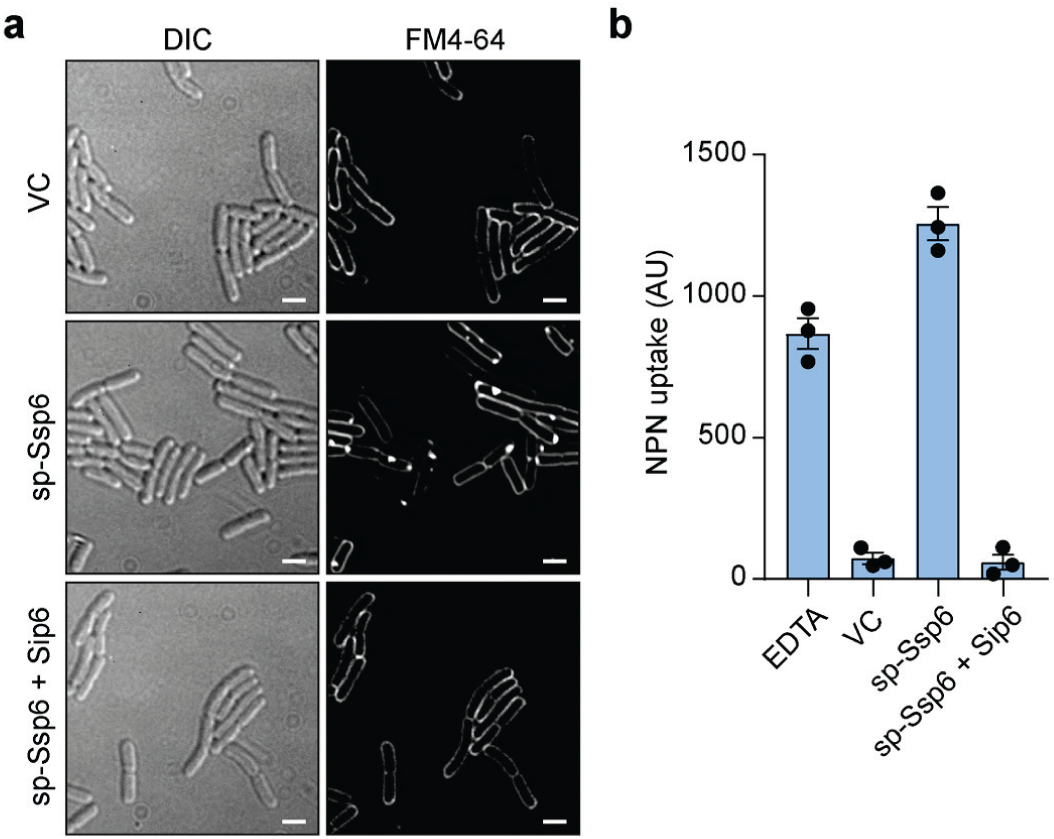
Ssp6 intoxication affects the outer membrane. (a) Visualisation of cells of *E. coli* MG1655 carrying empty vector control (VC, pBAD18-Kn) or plasmids directing the expression of Ssp6 fused with an N-terminal OmpA signal peptide (sp-Ssp6), either alone or with Sip6, using the membrane stain FM4-64 and fluorescence microscopy. FM4-64 staining was performed following growth in liquid LB containing 0.2% L-arabinose. Panels show DIC image (left) and FM4-64 channel (right). Scale bar 2 µm. Images are representative of four independent experiments. (b) Measurement of NPN uptake by *E. coli* expressing sp-Ssp6 alone or with Sip6, as in part a. NPN accumulation is expressed as arbitrary fluorescence units (AU) and bars show mean +/-SEM, with individual data points superimposed (n = 3 independent experiments).

### Ssp6 defines a new family of cation-selective pore-forming T6SS effectors occurring widely in the Enterobacteriaceae

Here we have shown that the anti-bacterial effector Ssp6 is a cation-selective pore-forming toxin. However it does not share sequence identity or predicted structural homology with previously-described T6SS effectors, including those proposed to be pore-forming effectors^12, 13^. In order to determine how widely Ssp6-like effectors occur in other organisms, we used HMMER homology searching to interrogate a database of complete, published bacterial genome sequences. Homologues of Ssp6 were found to be widespread across the family Enterobacteriaceae. We identified 95 homologues in 38 different species, with up to three Ssp6-like proteins encoded within the genomes of individual strains. For selected examples, the phylogenetic relationship of the identified Ssp6-like proteins and the genetic context of their encoding genes is depicted in Fig. 6, with the full set described in Supplementary Fig. 6 and Supplementary Table 3. In each case, a small open reading frame can be identified immediately downstream of the *ssp6*-like gene which is predicted to encode the corresponding immunity protein (Fig. 6). Whilst some of these share readily-detectable similarity with Sip6, due to the very small length of the proteins, we did not attempt further analysis of their phylogeny and relatedness. Examining the genetic context of the *ssp6*-like genes, we observed that this is variable between, and even within, species. In some cases, Ssp6-like effectors are encoded away from any other T6SS genes, as for Ssp6 itself. In other cases, Ssp6-like effectors are located within T6SS gene clusters, for example in *Enterobacter cloacae* EcWSU1, or with so-called ‘orphan’ Hcp genes, for example in some strains of *Klebsiella pneumoniae*, supporting their assignment as T6SS-dependent effector proteins (Fig. 6, Supplementary Fig. 6).

**Figure 6.**
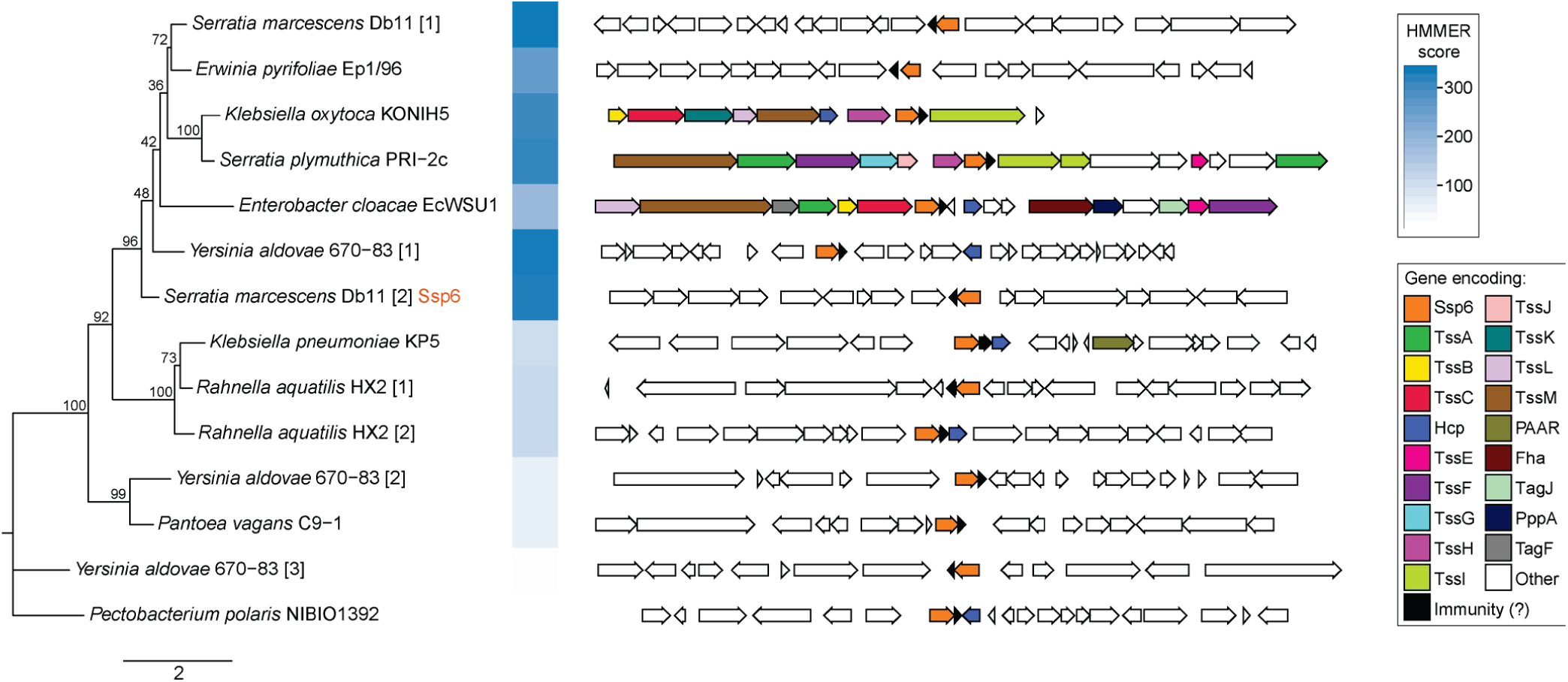
Genes encoding Ssp6-like effectors are widespread in Enterobacteriaceae and can be linked with Type VI secretion system genes. Phylogenetic tree of selected Ssp6 homologues identified using HMMER homology searching of complete bacterial genomic sequences (left) and the genetic context of the corresponding encoding gene (right). Where a particular organism encodes more than one Ssp6 homologue, each homologue is indicated by square brackets after the organism name. Bootstrap values are indicated on the tree and the scale indicates number of substitutions per site. In the genetic loci, conserved T6SS genes are coloured as per the legend; Ssp6 homologues are orange and genes encoding known or putative immunity proteins are black. The full set of identified homologues and details of the bacterial genome sequences can be found in Supplementary Figure 6 and Supplementary Table 3.

Interestingly, our bioinformatics analysis revealed that *S. marcescens* Db10 possesses a second homologue of Ssp6, SMDB11_0810, encoded elsewhere in the genome together with a homologue of Sip6, SMDB11_0809 (Fig. 6, Supplementary Fig. 7a). Thus, we tested whether these proteins represent a novel T6SS effector-immunity pair. We found that a target strain lacking the putative immunity protein (Δ*SMDB11_0810*-*0809*) does not display any reduction in recovery when co-cultured with a wild type attacking strain compared with a T6SS-inactive attacker. We also tested a target strain lacking both SMDB11_0809 and Sip6 (Δ*ssp6*Δ*sip6*, Δ*SMDB11_0810*-*0809*), but this strain showed no sensitivity to SMDB11_0810, only to Ssp6 (Supplementary Fig. 7b). This lack of SMDB11_0810 activity is most likely due to a lack of expression, since we did not detect this protein in the total cellular proteome of *S. marcescens* Db10 grown on solid LB media^16^ and have also never detected it in our secretome studies^14, 15^. Therefore we asked whether SMDB11_0810 displays any toxicity when artificially expressed and directed to the periplasm in *E. coli*. In this context, SMDB11_0810 induces detectable depolarisation in a small fraction of the cells. Whilst modest, this effect was recovered back to the level of the empty vector control by the co-expression of the putative immunity protein, SMDB11_0809 (Supplementary Fig. 7c). Expression of SMDB11_0810 also led to an increase NPN uptake and, therefore outer membrane permeability, albeit to a lesser extent than Ssp6. Again, co-expression of SMDB11_0809 was able to reverse the impact of SMDB11_0810, suggesting that SMDB11_0810 possesses at least residual anti-bacterial activity and SMDB11_0809 represents a functional cognate immunity protein.

## Discussion

In this study, we have determined the mode of action by which a novel T6SS effector, Ssp6, causes growth inhibition in intoxicated bacterial cells. We have demonstrated that Ssp6 acts by forming cation-specific channels, leading to inner membrane depolarisation and thus cell de-energisation, and that Ssp6 intoxication can also lead to increased outer membrane permeability. Importantly, Ssp6 is the founding member of a new family of T6SS-delivered, ion-selective pore-forming toxins, which are distinct from previously-described T6SS-effector proteins, including those proposed to form channels or pores.

To date, two T6SS-dependent effectors causing membrane depolarisation have been identified, VasX and Tse4^12, 13^. VasX displays some structural homology with pore-forming colicins and was shown to disrupt the membrane potential with simultaneous permeabilisation of the inner membrane^13, 26, 27^. Conversely, Tse4 disrupts the membrane potential without compromising the membrane permeability of intoxicated cells^12^. Thus, VasX is thought to form a large, non-selective pore, which would cause leakage of ions and other cellular contents^13, 27, 28^, whilst Tse4 was suggested to form a cation-selective pore^12^. Whilst we did not detect sequence or predicted structural similarity between Ssp6 and Tse4 or VasX, our data showed that Ssp6 can cause depolarisation of targeted cells without a corresponding increase in permeability of the inner membrane, suggesting that the Ssp6 and Tse4 modes of action may be similar. We speculate that the reason why a small fraction of the cells depolarised by Ssp6 also show an increase in membrane permeability is due to a downstream, secondary effect of being unable to maintain proper membrane or cell wall integrity.

Importantly, for the first time, we confirmed the ability of a T6SS effector, Ssp6, to form an ion-selective pore *in vitro* using artificial membranes. Ssp6 displayed a preference for monovalent cations compared with divalent cations, whilst surprisingly being impermeant to protons. The electron transport chain generates an electrochemical gradient of protons, the proton motive force (PMF), by extrusion of protons across the bacterial inner membrane^29^. The PMF is comprised of the transmembrane electrical potential, ΔΨ (negative inside the cell), and the transmembrane pH gradient, ΔpH^30, 31^. PMF is used to actively transport solutes against their electrochemical gradient, determining accumulation of K^+^ within the cell and extrusion of Na^+^ outside the cell^32, 33^. These processes contribute to generating an electrochemical gradient and maintaining the ΔΨ component of the PMF^29^. Based on the *in vitro* data, dissipation of the membrane potential through the action of Ssp6 is caused by an influx of cations into the cell. Whilst ion concentrations inside and outside bacterial cells can vary with growth and physiological conditions, in this study, co-culture assays measuring Ssp6 intoxication were performed on solid LB media containing 170 mM NaCl (with only residual K^+^, around 3-6 mM). Thus we speculate that the high concentration of Na^+^ outside of the cells, compared with the cytoplasm, will generate a Na^+^ electrochemical gradient that represents the driving force to cause influx of Na^+^ through Ssp6 pores, thereby dissipating the ΔΨ component of the PMF. Since Ssp6 pores were shown to be impermeant to H^+^, the ΔpH component would not be affected, as also reported for Tse4^12^.

Consistent with the ion-selective membrane depolarisation mechanism defined *in vitro*, intoxication by Ssp6 caused growth inhibition rather than cell lysis *in vivo*. In contrast, it has been reported that colicins, such as colicin E1 and N, which form a non-specific pore in the membrane of susceptible cells will ultimately cause lysis^34–36^. Nevertheless, depolarisation alone results in ATP depletion and general disruption of normal cell functions, such as Sec- and Tat-dependent protein export or solute and nutrient transport^29, 37–41^. Thus we hypothesize that whilst formation of non-selective pores allowing passage of large molecules could cause a more drastic leakage of cell contents and typically lead to lysis of intoxicated cells, formation of a cation-selective pore, such as Ssp6, has less drastic effects which are nevertheless sufficient to cause growth inhibition.

In general, pore-forming toxins (PFTs) can be classified into two large groups, α-PFTs and β-PFTs, based on whether their membrane spanning domain is composed of α-helices or β-barrels^42–44^. Whilst structural information will be required to fully understand the arrangement and mechanism of Ssp6 pores, secondary structure predictions indicate substantial α-helical content and thus likely an α-PFT classification. β-PFTs oligomerise at the membrane surface, forming an intermediate pre-pore which will then insert into the membrane upon reaching a threshold size^45, 46^. For α-PFTs, membrane insertion and oligomerisation are concomitant processes that can lead to formation of partially-assembled but active pores or complete pores^47, 48^. In both cases, oligomerisation and insertion in the membrane is observed when a critical concentration of monomers is reached^42^. In contrast with immunity proteins for colicin PFTs, which have been reported to be localised in the inner membrane^49, 50^, Sip6 is localised in the outer membrane. Localisation of Sip6 in the outer membrane may represent a means to sequester incoming Ssp6 away from the inner membrane and prevent free Ssp6 from reaching a critical concentration for pore formation. Its localisation might additionally reflect a second function in avoiding direct Ssp6 damage to the outer membrane. Apart from phospholipase effectors which are likely to be able to affect the inner leaflet of the outer membrane, to date there are no reports of T6SS-dependent effectors which directly damage the outer membrane. Whilst our data reveal that Ssp6 intoxication can increase outer membrane permeability and perhaps modify the distribution of its lipids, they do not distinguish between damage to the outer membrane being directly caused by Ssp6 and an indirect effect of Ssp6 downstream of inner membrane depolarisation.

Finally, we asked whether Ssp6 is unique to *Serratia marcescens* or if this type of T6SS effector occurs more widely. Analysis of whole-genome sequencing data revealed that homologues of Ssp6 are restricted to the Enterobacteriaceae but occur widely within this family, in at least 38 different species. This is likely to be an underestimate since our HHMER analysis was performed using only complete, published genome sequences, whereas initial analysis identified further homologues in, for example, clinical isolates of *E. coli* whose genomes were not fully sequenced. Ssp6-like effectors, like many anti-bacterial T6SS effectors, appear to be horizontally acquired, being present in only some strains of a given species and with variable genetic locations. In some cases, genes encoding Ssp6-like proteins are within a main T6SS gene cluster (encoding most or all of the structural and regulatory components making up the machinery) or in distant ‘orphan’ loci containing *hcp* genes. Genes encoding components of the expelled puncturing structure, Hcp, VgrG and PAAR proteins, are often present in multiple copies, with individual homologues required for delivery of particular effectors. In many cases, effectors are genetically linked with their cognate delivery protein^1^. Thus Ssp6-like proteins encoded adjacent to orphan *hcp* genes (or linked with an *hcp* gene in a T6SS gene cluster) are likely to be dependent on interaction with that Hcp homologue for their delivery. This is in agreement with previous findings consistent with Ssp6 being an Hcp-dependent effector in *S. marcescens* Db10^14^. One interesting example of *ssp6* context is in *Enterobacter cloacae* EcWSU1, which possesses a very similar T6SS gene cluster to *S. marcescens* Db10. In EcWSU1, the *ssp6*-like gene is present between *tssC* and *hcp* within this T6SS gene cluster, in the same position as the peptidoglycan hydrolase effector-immunity pair *ssp1-rap1a* in Db10, consistent with the idea of this being an effector acquisition/exchange hotspot^51^. This analysis also revealed the presence of a second Ssp6-Sip6-like pair of proteins encoded in the genome of *S. marcescens* Db10, SMDB11_0810 and _0809. Given that these are not expressed under our normal conditions, and show only limited activity when overexpressed in an equivalent manner to that in which we studied Ssp6, we speculate that they might be expressed and show higher efficacy under quite different physiological or environmental conditions. The idea of conditional effector efficacy has been supported by studies in *P. aeruginosa*^12^ and the fact that several effectors of the same family are often observed in the same organism may support the idea that two related effectors with different regulation and/or specificity could provide a bet-hedging strategy to deal with different conditions and competitors. For example, Db10 itself possesses two related Tae4-family peptidoglycan hydrolase effectors with distinct substrate specificity *in vitro* and different efficacy *in vivo*^51, 52^.

In conclusion, this study has revealed that Ssp6 is the founder member of a new family of T6SS-dependent, cation-selective pore-forming anti-bacterial effectors. This toxic activity leads to depolarisation of the inner membrane, disruption of outer membrane integrity and, consequently, to inhibition of growth in targeted cells. We propose that this family could be named Tpe1, as the first example of a <u>T</u>6SS-dependent <u>p</u>ore-forming <u>e</u>ffector whose activity has been confirmed *in vitro*. At the molecular level, it will be of great interest to determine how these proteins are able to generate a gated pore with such defined ion selectivity, allowing mono- and di-valent metal cations, but not protons, to pass through a membrane. At the population level, Ssp6-family effectors further expand the already-impressive repertoire of toxins bacteria can deploy to compete with each other. Elucidating the basis of competitive inter-bacterial interactions is vital to understand and utilise their capacity to shape the composition and dynamics of diverse polymicrobial communities, including those important for health, disease and biotechnological applications.

## Material and Methods

### Bacterial strains and plasmids

Bacterial strains and plasmids used in this study are detailed in Supplementary Table 1. Strains of *S. marcescens* Db10 carrying in-frame deletions or encoding epitope-tagged fusion proteins at the normal chromosomal location were generated by allelic exchange using the plasmid pKNG101^53^. Streptomycin-resistant derivatives were generated by transduction of the resistance gene from *S. marcescens* Db11, as previously described^51^. Primer sequences and details of construction are provided in Supplementary Table 2. Plasmids for constitutive expression of genes *in trans* were derived from pSUPROM while plasmids for arabinose-inducible expression were derived from pBAD18-Kn. Strains of *S. marcescens* were grown at 30°C on LB agar (LBA, 10 g/L tryptone, 5 g/L yeast extract, 10 g/L NaCl, 12 g/L agar), in liquid LB (10 g/L tryptone, 5 g/L yeast extract, 5 g/L NaCl) or in minimal media (40 mM K_2_HPO_4_, 15 mM KH_2_PO_4_, 0.1% [NH_4_]_2_SO_4_, 0.4 mM MgSO_4_, 0.2% [w/v] glucose), whilst for *E. coli* growth was at 37°C. When required, media were supplemented with kanamycin (Kn) 100 µg/ml, streptomycin (Sm) 100 µg/ml, or ampicillin (Ap) 100 µg/ml.

### Immunodetection of cellular and secreted proteins

Detection of Hcp in cellular and secreted fractions of cultures grown for 5 h in LB was performed as described^51^. For detection of Ssp6-HA, bacterial strains were grown for 5 h in LB. Cellular protein samples were prepared by resuspending cells recovered from 200 μL of culture in 100 μL of 2x SDS sample buffer (100 mM Tris-HCl [pH 6.8], 3.2% SDS, 3.2 mM EDTA, 16% glycerol, 0.2 mg/mL bromophenol blue, 2.5% β-mercaptoethanol). Secreted protein samples were prepared by precipitation from 15 ml of culture supernatant and resuspension in a final volume of 40 μL, according to the method described previously^51^. Ssp6-HA was detected using anti-HA primary antibody (MRC PPU Reagents, University of Dundee, 1:6,000) and HRP-conjugated anti-mouse secondary (1:10,000; Bio-Rad).

### Co-culture assays for T6SS-mediated antibacterial activity

T6SS-mediated anti-bacterial activity of strains of *S. marcescens* Db10 was measured as described^51, 53^. In brief, attacker and target strains were normalised to OD_600_ 0.5, mixed at a 1:1 ratio and co-cultured on solid LB at 30°C for 4 h unless stated otherwise. The number of surviving target cells was enumerated by serial dilution and viable counts on Sm-supplemented LB agar, with all target strains being Sm-resistant derivatives of the wild type or mutant of interest.

### Growth inhibition upon heterologous toxin expression

To study the impact of heterologous expression of Ssp6 from pBAD18-Kn-derived plasmids, freshly-transformed cells of *E. coli* MG1655 grown overnight on solid media were adjusted to OD_600_ of 1, serially diluted and 5 μl spotted onto LB plates containing either 0.2% D-glucose or 0.2% L-arabinose. For growth rate measurement in liquid culture, cultures were inoculated to a starting OD_600nm_ 0.02 in a volume of 25 mL LB and incubated at 30 °C at 200 rpm. Optical density at 600 nm (OD_600_) readings were acquired every 1 h. Expression from pBAD18-Kn-derived plasmids was induced with 0.2% L-arabinose at OD_600_ 0.2.

### Immunoprecipitation of Ssp6-HA and Sip6-FLAG

Cultures of *S. marcescens* strains carrying chromosomally-encoded Ssp6 with a C-terminal HA tag (Ssp6-HA) and/or Sip6 with a C-terminal triple FLAG tag (Sip6-FLAG) were grown in LB for 5 h to an OD_600_ of ∼4.5. Cells were recovered by centrifugation at 4000 *g* for 20 min, resuspended in 50 mM Tris-HCl pH 7.8 and lysed using an EmulsiFlex-C3 homogenizer (Avestin). Cell debris were removed by centrifugation (14,000 *g*, 45 min, 4°C) and 1 ml of lysate (corresponding to 50 ml of the original culture) was transferred into tubes containing 50 μl of pre-washed (3x) magnetic α-HA beads (NEB) and incubated for 2 h at 4°C, 20 rpm. The beads were then washed with 4 × 1 ml of wash buffer (20 mM Tris-HCl pH 7.8, 100 mM NaCl. 0.1% Triton X-100) and bound proteins were eluted by addition of 40 μl of 2x SDS sample buffer.

### Inner and outer membrane fractionation

Cultures of *S. marcescen*s were grown in 25 mL LB for 5 h to an OD_600_ of approximately 3. Cells were recovered by centrifugation at 4000 *g*, 4°C for 10 min and resuspended in 1 mL of 50 mM Tris-HCl pH 8. Cells were then lysed by sonication, unbroken cells removed by centrifugation (14.000g, 20 min, 4°C) and the cell-free lysate subjected to ultracentrifugation (200,000 *g*, 30 min, 4°C). The resulting supernatant was collected and represented the cytoplasm fraction. The pellet was resuspended in 1 mL of 50 mM Tris-HCl pH 8 and a small aliquot removed, representing the total membrane fraction. The detergent C8POE (octyl-poly-oxyethylenoxide) (Bachem) was then added to a final concentration of 2%, incubated at 37°C for 30 min and a second ultracentrifugation step performed (200,000 *g*, 30 min, 4°C). The resulting supernatant was collected, representing the outer membrane fraction in *S. marcescens*, while the pellet was resuspended in 1 mL of 50 mM Tris-HCl pH 8, representing the *S. marcescens* inner membrane fraction. Equal volumes of each fraction (corresponding to material from the equivalent number of cells) were separated by SDS-PAGE and subjected to immunoblotting. Anti-FLAG antibody (Sigma) for detection of Sip-FLAG was used at, 1:10,000, while anti-EFTu (Hycult Biotech) was used 1:20,000, both with horseradish peroxidase (HRP)-conjugated anti-mouse secondary antibody (1:10,000; Bio-Rad). Anti-OmpA antibody^54^ was used at 1:20,000 and a custom anti-TssL antibody (Eurogentec; see Supplementary Fig. 8) was used at 1:6,000, both with peroxidase-conjugated anti-rabbit secondary antibody (Bio-Rad, 1:10,000).

### Membrane potential and membrane permeability analysis

For analysis of co-cultures, attacker and target strains of *S. marcescens* Db10 were co-cultured on solid LB media for 4 h at 30°C, then cells were recovered and suspended in 1x PBS at 10^6^ cells/ mL. DiBAC_4_(3) (Bis-[1,3-Dibutylbarbituric Acid] Trimethine Oxonol; Thermo) at 10 µM final concentration and propidium iodide at 1 µM final concentration were added simultaneously to each cell suspension, followed by incubation in the dark for 30 min. For analysis of plasmid-based expression of sp-Ssp6, cultures of *E. coli* MG1655 carrying pBAD18-Kn derived plasmids were inoculated to a starting OD_600_ of 0.02 in 25 mL LB and incubated at 37°C for 1.5 h, then induced by the addition of 0.2% L-arabinose and grown for a further 1 h. Cells were then recovered and resuspended in 1x PBS and DiBAC_4_(3) and propidium iodide staining was performed as above. As a control, cells were treated with polymyxin B (300 µg/mL for *S. marcescens* and 2 µg/mL for *E. coli*) at 37 °C for 30 min prior to staining. Following staining with DiBAC_4_(3) and propidium iodide, cells were directly analysed in a FACS LRS Fortessa equipped with 488 nm and 561 nm lasers (Becton Dickinson), using thresholds on side and forward scatter to exclude electronic noise. Channels used were Alexa 488 (Ex 488 nm, Em 530/30nm) for DiBAC_4_(3) and Alexa 568 (Ex 561 nm, Em 610/20 nm) for propidium iodide. All bacterial suspensions were normalised to 10^6^ cells/ mL prior to analysis. Analysis was performed using FlowJo v10.4.2 (Treestar Inc.); example plots can be seen in Supplementary Fig. 2.

### NPN uptake assay

Cultures of *E. coli* MG1655 carrying pBAD18-Kn derived plasmids were inoculated to a starting OD_600_ of 0.02 in 25 mL LB and incubated at 37 °C for 1.5 h, then induced by the addition of 0.2% L-arabinose and grown for a further 1 h. Cells were normalised to OD_600_ 0.7, washed twice with NPN assay buffer (5 mM HEPES pH 7.2, 5 mM glucose) and resuspended in NPN assay buffer to a final OD_600_ of 1.4 as previously described^24, 25^. The assay was prepared in a 96-well optical-bottom black plate (Thermo), by addition of 20 µM NPN (1-N-phenylnapthylamine, Sigma) in a final volume of 200 µL. EDTA was added to a final concentration of 10 mM as a control for outer membrane permeabilization. Fluorescence changes were monitored using a Clariostar Monochromator Microplate reader (BMG Labtech), with a wavelength of 355 nm for excitation and 405 nm for emission. NPN uptake was calculated, as previously described^25^, with the following formula:

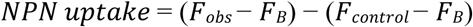

where F_*obs*_ represents the NPN fluorescence observed with *E. coli* strains carrying the test plasmids, F_B_ is the fluorescence in the absence of bacterial cells and F_control_ is the fluorescence with *E. coli* cells lacking a plasmid.

### FM-64 staining

For membrane staining using FM4-64 (Thermo), cultures of *E. coli* MG1655 carrying pBAD18-Kn derived plasmids were inoculated to a starting OD_600_ of 0.02 in 25 mL LB and incubated at 37°C for 1.5 h. When the exponential phase was reached (OD_600_ of 0.15-0.2), gene expression was induced by the addition of 0.2% L-arabinose followed by a further 1 h incubation. FM4-64 was added to a final concentration of 1 µM and samples were incubated at 37°C for 20 min. Three µL of stained cells were placed on a microscope slide layered with a pad of M9 media^55^ solidified by the addition of 1.5% agarose. Imaging was performed as described in the following section.

### Fluorescence Microscopy

For time-lapse experiments (Figure 2), cells of *S. marcescens* were pre-grown for 4 h in liquid minimal media to an OD_600_ of ∼0.3-0.4. Cultures were then normalized to OD_600_ 0.2, mixed in a ratio of 3:1 attacker:target, and 2 µL of the mixture placed on a pad of minimal media solidified by the addition of 1.5% UltraPure agarose (Invitrogen). Fluorescence imaging was performed using a DeltaVision Core widefield microscope (Applied Precision) mounted on an Olympus IX71 inverted stand with an Olympus 100X 1.4 NA lens and Cascade2_512 EMCCD camera with differential interference contrast (DIC) and fluorescence optics. Images were acquired with the following parameters: 512 × 512 pixels, 2-by-2 binning, with 11 Z sections spaced by 0.2 µm. GFP (target cells) and mCherry (attacker cells) were detected using a GFP filter set (Ex 485/20 nm, Em 530/25 nm) and mCherry filter set (Ex 542/82 nm, Em 603/78 nm), respectively. Independent fields of view were located and their XYZ positions were stored in order to capture images of the same coordinates every 30 min for 3 hours. Images were manually corrected for drift and, where necessary, adjusted for any loss of fluorescence during the timecourse.

Post-acquisition, images were deconvolved using softWoRx and stored and processed using OMERO software (http://openmicroscopy.org)^56^. Quantification of attacker and target cells was done using OMERO.mtools (http://www.openmicroscopy.org/site/products/partner/omero.mtools). All attacker strains carried cytoplasmic mCherry while target strains carried cytoplasmic GFP (Supplementary Table 1). For time-lapse experiments, microcolonies were chosen for analysis on the basis that they contained attacker and target cells which were in direct contact to allow T6SS-mediated attacks. The number of attacker and target cells in the microcolonies were counted at time point 0 h and 3 h. At least 70 cells (at t = 0 h) per strain per replicate were counted for the attackers and 19 cells per replicate were counted for the targets.

To quantify the relative number of attacker and target cells following co-culture under the conditions used for membrane potential and permeability staining (co-culture of attacker and target strains on solid LB media for 4 h at a starting ratio of 1:1, Fig. 3a), cells were recovered at the end of the co-culture, resuspended in LB and imaged as described above. Between 1450 and 1600 total cells (mixed target and attackers) per replicate for each condition were counted.

In the case of FM4-64 staining, imaging was performed using a CoolSnap HQ2 camera (Photometrics), with differential interference contrast (DIC) and fluorescence optics. Images were acquired with the following parameters: 512 × 512 pixels, 1-by-1 binning, with 11 Z sections spaced by 0.2 µm. FM4-64 fluorescence was detected using a TRITC (tetramethylrhodamine) filter set (Ex 540/25 nm, Em 605/55 nm). Images were adjusted for clear visualisation of the cell outline in each case.

### Purification of MBP-Ssp6

For purification of MBP-Ssp6, *E. coli* C43 (DE3) cells transformed with a pNIFTY-MBP-derived plasmid encoding Ssp6 fused with N-terminal MBP were inoculated to a starting OD_600_ 0.05 in 4 L LB, grown at 30°C, 200 rpm for 3 h, induced with 0.5 mM IPTG and then incubated for 16 h at 16°C. Cells were recovered by centrifugation (4000 *g*, 30 min), resuspended in 40 ml of Buffer A (50 mM Tris-HCl pH 8, 500 mM NaCl) in presence of cOmplete™ EDTA-free protease inhibitor (Sigma) and lysed using an EmulsiFlex-C3 homogenizer (Avestin). The lysate was cleared by centrifugation (14,000 *g*, 45 min, 4°C), filtered through a 0.45 µm filter, and loaded onto 1 mL MBP Trap™ HP column (GE Healthcare) following equilibration with Buffer A. Elution was achieved using 10 column volumes of Buffer B (50 mM Tris-HCl pH 8, 500 mM NaCl, 10 mM maltose). The eluted fraction was separated by size exclusion chromatography using a Superose 6 Increase 10/300 column (GE Healthcare) and a buffer containing 50 mM Tris-HCl pH 8, 150 mM NaCl, 10% glycerol.

### Electrophysiology measurements and analysis

Planar lipid bilayers were prepared as described previously^57^. Briefly, bovine phosphatidylethanolamine lipids (Stratech Scientific Ltd) were resuspended in decane at a final concentration of 30 mg/mL. Planar phospholipid bilayers were formed across a 150 µm diameter aperture in a partition that separates two 1 mL compartments, the *cis* and the *trans* chambers. MBP-Ssp6 was added to the *cis* chamber. The *trans* chamber was held at 0 mV (ground potential) while the *cis* chamber was clamped at different holding potentials relative to ground. The transmembrane current was measured under voltage clamp conditions using a BC-525C amplifier (Warner Instruments, Harvard Instruments). Channel recordings were low-pass filtered at 10 kHz with a four-pole Bessel filter, digitized at 100 kHz using a National Instruments acquisition interfact interface (NIDAQ-MX, National Instruments, Austin, TX) and recorded on a computer hard drive using WindEDR 3.05 Software (John Dempster, University of Strathclyde, Glasgow, UK). Current fluctuations were measured over ≤ 30s and recordings were then filtered with WinEDR 3.8.6 low pass digital filter at 800 Hz (−3dB) using a low pass digital filter implemented in WinEDR 3.05. For experiments in which a nonsymmetrical KCl gradient was used, the KCl solution in the *cis* chamber contained 510 mM KCl, 10 mM HEPES pH 7.2, while the KCl solution in the *trans* chamber contained 210 mM KCl, 10 mM HEPES pH 7.2. For bi-ionic relative permeability studies, the KCl solution contained 210 mM KCl, 10 mM HEPES pH 7.2; the NaCl solution contained 210 mM NaCl, 10 mM HEPES pH 7.2; and the CaCl_2_ solution contained 210 mM CaCl_2_, 10 mM HEPES pH 7.2.

Noise analysis was performed by subdividing current recordings into segments in time, with each segment containing N samples. For each holding potential, current fluctuations were measured over 20-30 s. For each segment, the mean current was plotted against time and computed using the following formula:

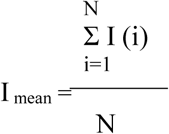

Where I(i) is the amplitude of the ith current and N is the number of the samples in the analysed segment. The mean data obtained from multiple replicates were subsequently plotted as a function of voltage. Predicted reversal potentials were calculated using the Nernst equation.

The relative permeability ratio when comparing relative permeability of monovalent cations was calculated using the Goldman-Hodgkin-Katz equation^58, 59^:

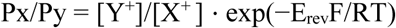

Where R is the ideal gas constant (8.314 J mol^−1^), T is the temperature expressed in kelvin, F is the Faraday constant (9.6485 × 10^4^ C mol^−1^) and E_rev_ is the reversal potential, which was taken to be the holding potential at which transmembrane current fluctuations were at a minimum.

The relative monovalent to divalent cations permeability ratio was calculated using the Fatt-Ginsborg equation^60^

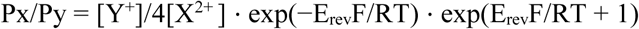

Junction potentials were calculated using Clampex software version 10.2 (Molecular Devices) and subtracted from the reversal potential obtained for each experiment.

### Identification of Ssp6-like effector family

#### 1. Selection of genome sequences

Genome sequences were downloaded from the RefSeq database^61^ (ftp://ftp.ncbi.nlm.nih.gov/genomes/ASSEMBLY_REPORTS/assembly_summary_refseq.txt, as of 4th Feb 2019). Genome sequences that were designated “Complete Genomes” and had previously been published were selected for analysis. Genomes were annotated using prokka (v1.13.3)^62^ and CDS protein sequences extracted using a custom script (gff_to_faa.py). Genomes were selected for further analysis based on possession of both at least one gene encoding an Ssp6-like protein, and at least one T6SS-encoding gene cluster. To achieve these criteria, two steps were performed. First, hmmsearch from the HMMER suite (v 3.1b2)^63^ was used to identify genomes containing genes encoding Ssp6-like proteins, based on a small, manually-curated alignment of non-redundant Ssp6 homologues which had originally been identified using BLASTp; the cutoff value was a HMMER bit score of 20 over the overall sequence/profile comparison. Second, T6SS-encoding gene clusters were identified as a locus containing at least 9 T6SS component genes (components identified using HMMER, cutoff value as above) in a contiguous set with no more than 8 unrelated genes between each known T6SS gene, performed using a custom script (hamburger.py). HMM models were taken from PFAM, or created from protein sequences stored in the Secret6 database (using hmmbuild) for accessory and core components of the T6SS respectively. All models and the alignments they are based on (if applicable) are stored in https://github.com/djw533/ssp6-paper/tree/master/models. All custom scripts can be found at https://github.com/djw533/ssp6-paper/tree/master/scripts.

#### 2. Extraction and analysis of Ssp6-like protein sequences and genetic loci

Ssp6-encoding loci of approximately 20K nucleotides (10 Kb upstream and downstream of the Ssp6 HMMER hit) were extracted using the script hamburger.py from genome sequences that satisfied the requirements stated in part 1. Extracted loci were then subsequently analysed for possession of T6SS related genes using hamburger.py a second time, as in part 1. Ssp6-like protein sequences were aligned using MUSCLE (v3.8.31)^64^ and trees drawn using IQTREE (v1.6.5)^65^ with 1000 ultrafast bootstraps. Trees were visualised using the R package ggtree (v1.15.6)^66^, and associated genomic context depicted using ggplot2 (v3.1.1)^67^ and gggenes (v0.3.2) (https://cran.r-project.org/web/packages/gggenes/). An R script https://github.com/djw533/ssp6-paper/blob/master/scripts/Plotting_ssp6_figures.R) was used to plot figures from the above results (stored at https://github.com/djw533/ssp6-paper/tree/master/results).

### Data availability statement

All data supporting the findings of this study are available within the paper and its supplementary information files.

## Supporting information

Combined Supplementary Information

## Acknowledgements

This work was supported by the Wellcome Trust (104556/Z/14/Z, Senior Fellowship in Basic Biomedical Science to S.J.C.; 097818/Z/11/B and 109118/Z/15/Z, PhD studentships to University of Dundee), the MRC (MR/K000111X/1, New Investigator Research Grant to S.J.C.) and the Royal Society of Edinburgh (Biomedical Personal Research Fellowship to S.J.P.). We thank Roland Freudl for the gift of anti-OmpA antibody; Adam Ostrowski for construction of strains AO07 and AO08; Gal Horesh, Amy Dorward and Gavin Robertson for expert assistance; the Flow Cytometry and Cell Sorting Facility at the University of Dundee; and the Dundee Imaging Facility (supported by Wellcome Trust [097945/B/11/Z] and MRC [MR/K015869/1]) awards).

## Author Contributions

G.M., K.T. and S.J.C. conceived the study; G.M., K.T., H.S., S.J.P. and S.J.C. designed experiments and analysed data; G.M., K.T. and L.M. performed experimental work; D.J.W. performed bioinformatics analyses; G.M. and S.J.C wrote the manuscript with input from all the other authors.

