## Supplementary material for "A new family of Type VI secretion system-delivered effector proteins displays ion-selective pore-forming activity": Combined Supplementary Information

#### Supplementary Figure 1

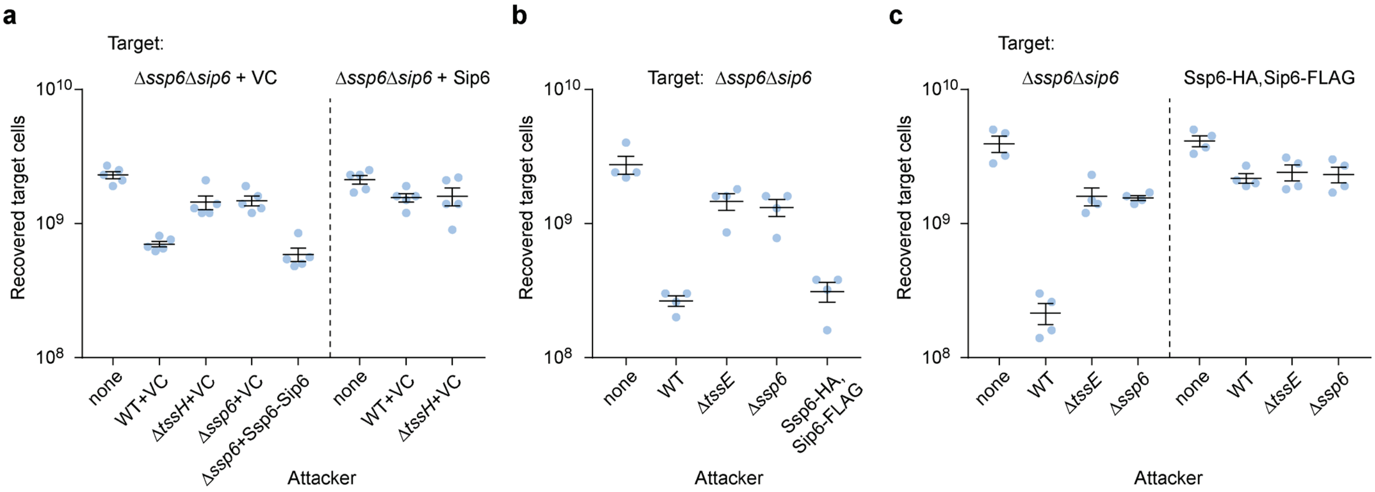


**Supplementary Figure 1. Complementation and functionality of strains carrying Ssp6 and Sip6 deletions or epitope tag fusions.** (a) Recovery of target strain *S. marcescens* Db10 Δ*ssp6*Δ*sip6* carrying the empty vector control (+VC, pSUPROM) or a plasmid directing the expression of Sip6-HA (+Sip6), following 7.5 h co-culture with wild type (WT) or mutant strains (Δ*tssH*, Δ*ssp6*) of Db10 carrying the empty vector control (+VC, pBAD18-Kan) or a plasmid directing the expression of Ssp6 and Sip6 (+Ssp6-Sip6). Expression of Ssp6 and Sip6 was induced by addition of 0.02% L-arabinose; Sip6 was included in the Ssp6 construct to prevent toxicity from overexpression of Ssp6. (b) Recovery of the Δ*ssp6*Δ*sip6* mutant following co-culture with wild type *S. marcescens* Db10, the Δ*tssE* mutant and the strain encoding Ssp6-HA and Sip6-FLAG fusion proteins at the normal chromosomal location. (c) Recovery of the wild type and the strain encoding Ssp6-HA and Sip6-FLAG fusion proteins as targets, when co-cultured with wild type, Δ*tssE* or Δ*ssp6* attackers. All target strains used in the study represent streptomycin-resistant derivatives of the strain indicated. In all parts, individual data points are overlaid with the mean +/- SEM (n=4 biological replicates, or n=5 in part a); none, target cells incubated with sterile media alone.

### Supplementary Figure 2

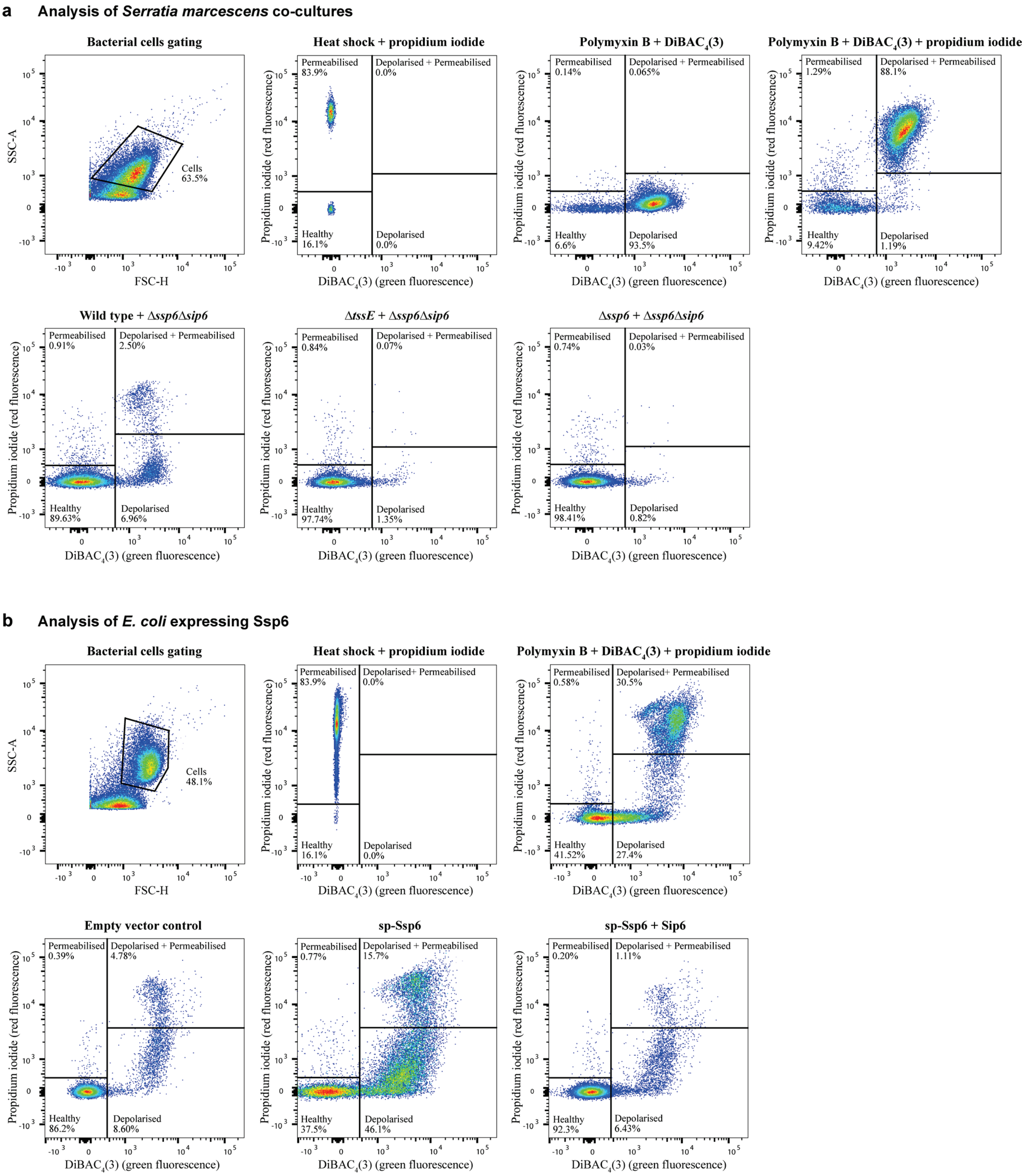


**Supplementary Figure 2. Examples of individual flow cytometry experiments from the analyses presented in Figure 3.** (a) Analysis of *S. marcescens* co-culture experiments, as Figure 3a. Top panels illustrate the gating strategy. Bacterial cells were selected using side scatter (SSC-A) vs. forward scatter (FSC-H). Cells subjected to heat shock treatment and stained with propidium iodide only were used to define the permeabilized cells quadrant (red fluorescence), whilst treatment with 300 μg/mL polymyxin B and staining with DiBAC$_4$(3) only was used to define the depolarised cells quadrant (green fluorescence). Cells treated with polymyxin B and stained with DiBAC$_4$(3) and propidium iodide were used to establish the quadrant for cells that are simultaneously depolarised and permeabilized (green and red fluorescence). Bottom panels show representative plots from experiments involving co-culture of the Ssp6-susceptible target strain Δ*ssp6*Δ*sip6* with wild type (WT) or control (Δ*ssp6* or Δ*tssE*) attacker strains of *S. marcescens* Db10, followed by simultaneous staining with propidium iodide and DiBAC$_4$(3). (b) Analysis of *E. coli* cells expressing sp-Ssp6, as Figure 3b. Top panels illustrate the gating strategy. Bacterial cells were selected using SSC-A vs. FSC-H. Cells subjected to heat shock treatment and stained with propidium iodide only were used to define the permeabilized cells quadrant (red fluorescence), whilst treatment with 2 μg/mL polymyxin B and staining with both propidium iodide and DiBAC$_4$(3) defined the quadrant of cells that are simultaneously depolarised and permeabilized (green and red fluorescence) and determined the separation from the quadrant containing cells that are depolarised but not permeabilised (green fluorescence). Bottom panels show representative plots from experiments analysing *E. coli* cells expressing sp-Ssp6 or sp-Ssp6 + Sip6, or healthy control cells carrying the empty vector (VC).

#### Supplementary Figure 3

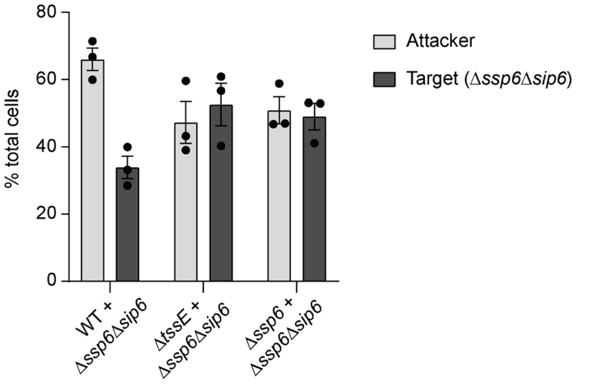


**Supplementary Figure 3. Quantification of the relative proportion of attacker and target cells following co-culture under the conditions used for membrane potential and permeability analysis in Figure 3a.** The Ssp6-susceptible target strain, Δ*ssp6*Δ*sip6* expressing cytoplasmic GFP, was co-cultured with wild type (WT) or mutant (Δ*ssp6* or Δ*tssE*) attacker strains expressing cytoplasmic mCherry for 4 h at an initial ratio of 1:1. Cells were recovered, imaged by fluorescence microscopy and numbers of green (GFP) and red (mCherry) cells counted using OMERO mTools. The relative numbers of attacker and target cells for each condition following co-culture is represented as a percentage of the total cell population. Around 1500 cells in total were counted for each condition in each replicate. Bars show mean +/- SEM, with individual data points superimposed (n=3 independent experiments).

#### Supplementary Figure 4

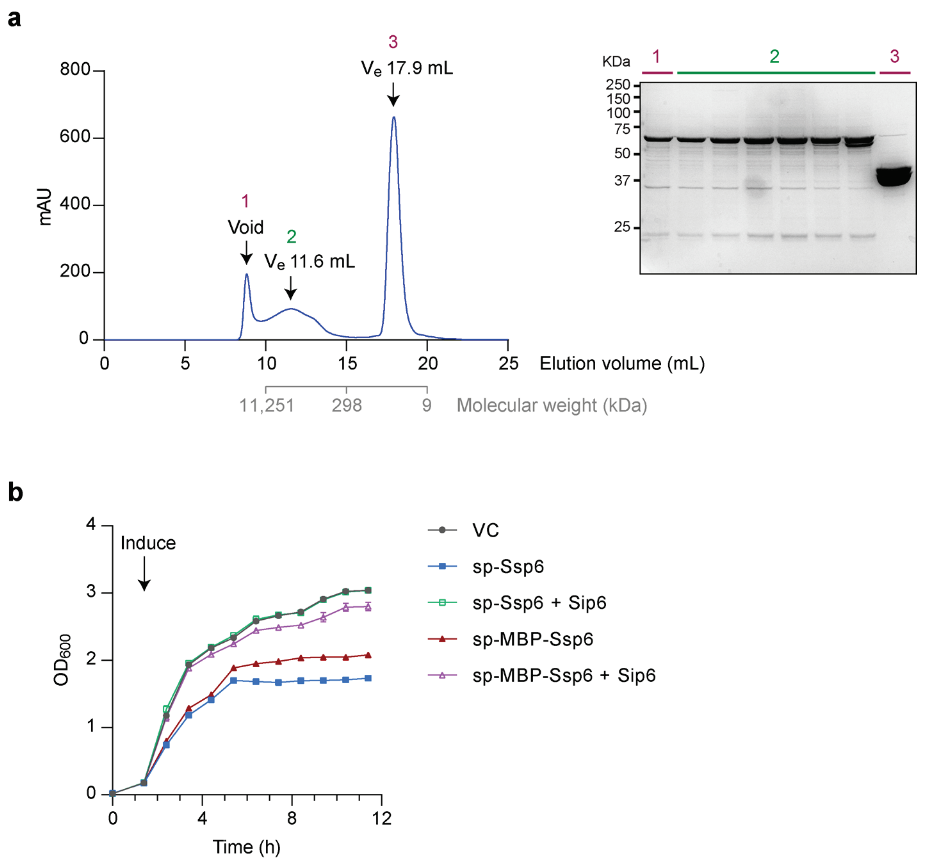


**Supplementary Figure 4. MBP-Ssp6 is oligomeric and retains toxin function.** (a) Size exclusion chromatography of MBP-Ssp6, following its initial isolation by dextrin affinity purification, using a calibrated Superose 6 Increase 10/300 GL column (GE Healthcare). For each peak, the elution volume ($V_e$) is indicated. Analysis of fractions eluted during size exclusion chromatography by SDS-PAGE and Coomassie staining is shown on the right, with peak 2 (green fractions) containing the MBP-Ssp6 used for subsequent analyses. Peak 3 contains MBP only. (b) Growth of *E. coli* MG1655 carrying empty vector (VC, pBAD18-Kn) or plasmids directing the expression of the MBP-Ssp6 fusion protein with an N-terminal OmpA signal peptide (sp-MBP-Ssp6) or native Ssp6 with signal peptide (sp-Ssp6), each with or without co-expression of Sip6 (+ Sip6), in LB containing 0.2% L-arabinose. Point show mean ± SEM (n=3 biological replicates).

**Supplementary Figure 5**

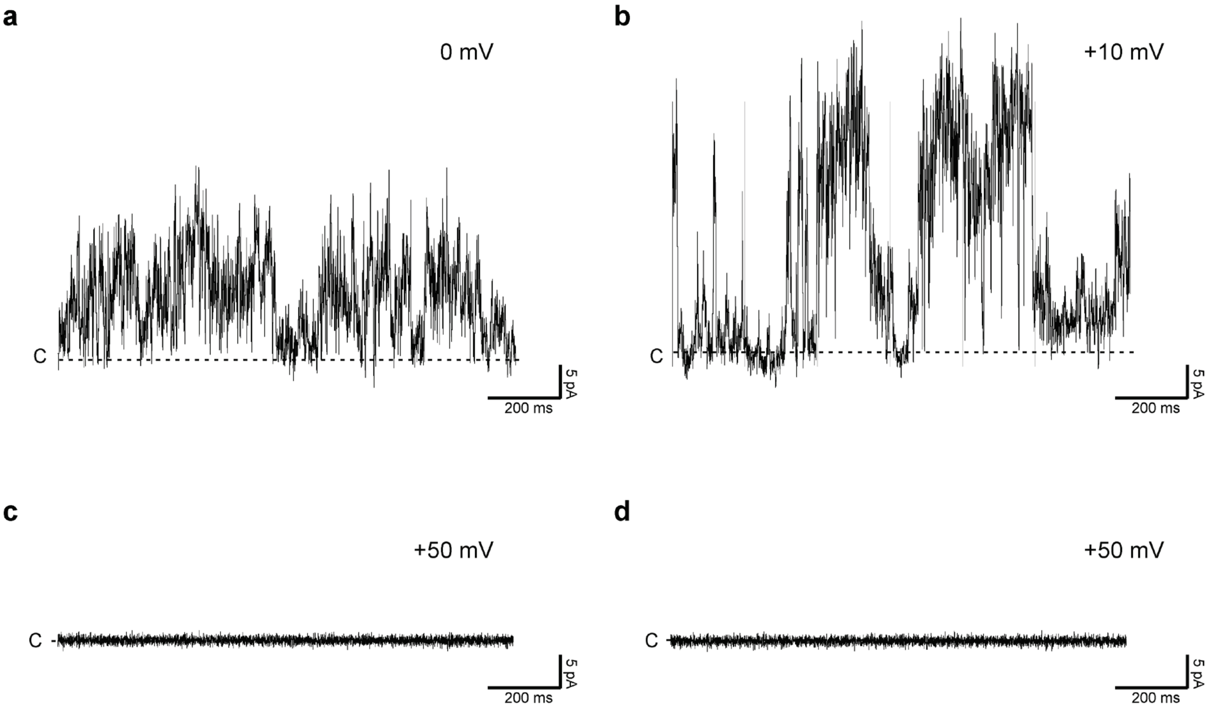


**Supplementary Figure 5. Electrophysiology analysis of Ssp6.** (a,b) Examples of single current fluctuations from MBP-Ssp6 in a KCl gradient showing that the pore generated by Ssp6-MBP exists in different states and one to multiple copies can be incorporated into a lipid bilayer, causing great variability of the mean current measured. (c,d) Measurement of current fluctuations following addition of purification buffer (part c) or MBP alone (part d). The traces shown are representative of three independent experiments. Throughout, the dotted line indicates the closed (C) state of the pore.

### Supplementary Figure 6

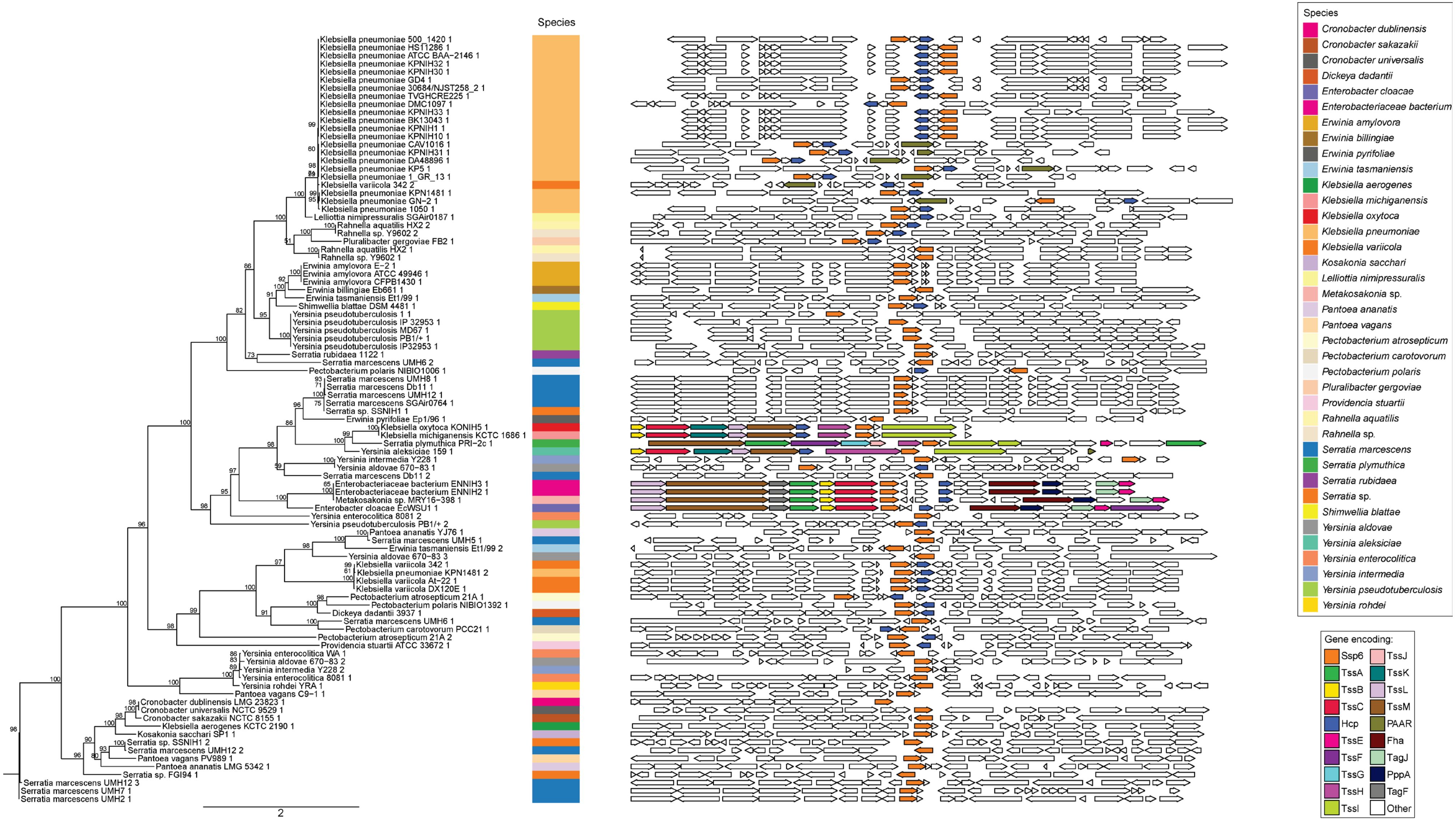

**Supplementary Figure 6. Full set of Ssp6-like proteins identified through bioinformatics analysis.** Phylogenetic tree and genetic context of homologues of Ssp6 identified using a HMMER search of complete, published bacterial genome sequences. Representative examples taken from this set are depicted in Figure 6 and details of the genome sequences and identified homologues are given in Supplementary Table 3. Note that small open reading frames downstream of *ssp6* encoding putative immunity proteins are frequently missed by the automated annotation; hence absence of such an open reading frame in the schematic does not imply such an immunity does not exist. Manual examination of a selection of such cases revealed that a candidate open reading frame downstream of the *ssp6*-like gene was present in each case as predicted. Genes whose boundaries extend beyond the selected region of interest are also not depicted.

#### Supplementary Figure 7

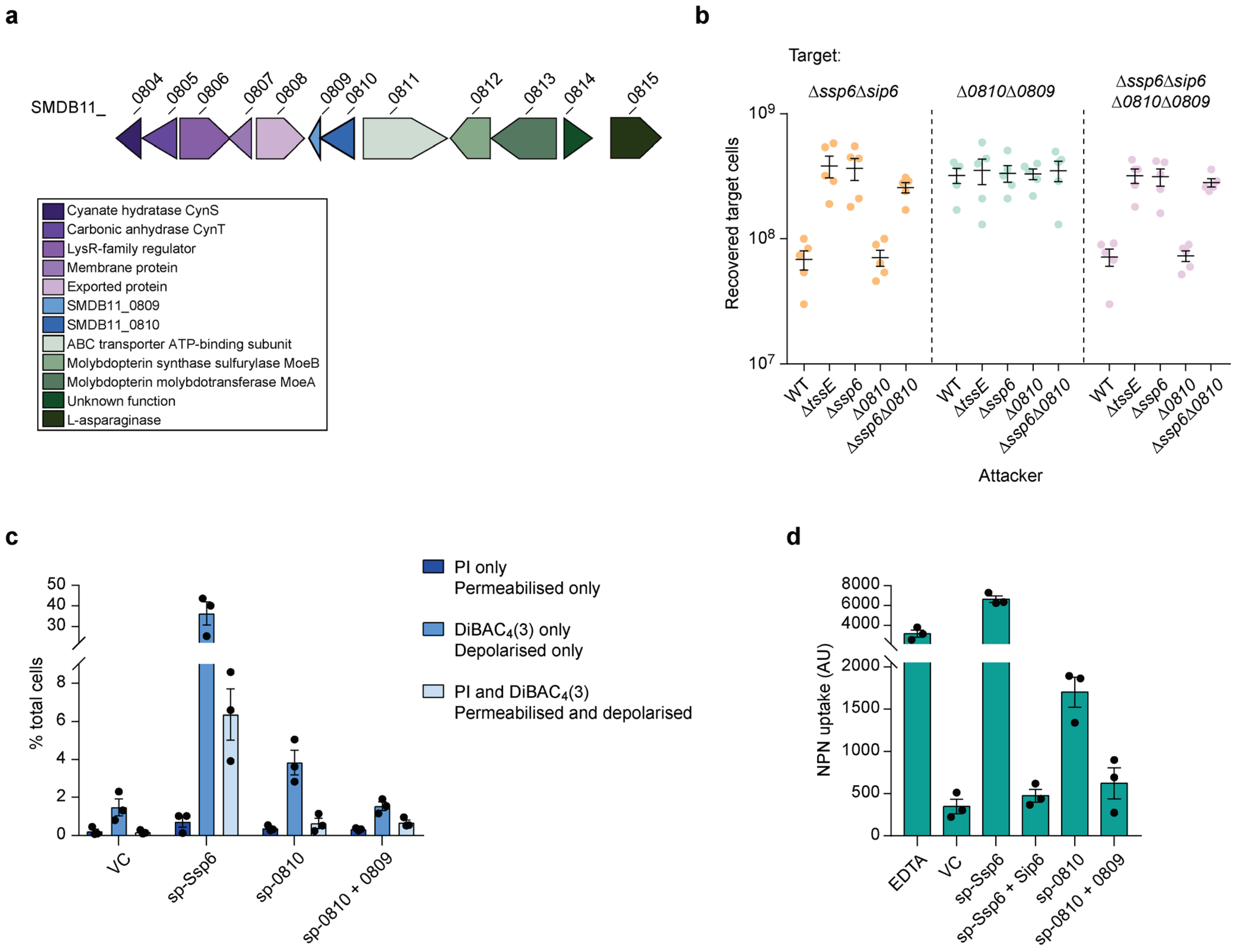


**Supplementary Figure 7. SMDB11_0810 and SMDB11_0809 represent 'silent' homologues of the Ssp6-Sip6 effector-immunity pair.** (a) Schematic representation of the genomic context of SMDB11_0810 and SMDB11_0809, with genomic identifiers (SMDB11_xxxx) provided above each gene and predicted protein functions in the box below. (b) Recovery of target strains of *S. marcescens* Db10, Δ*ssp6*Δ*sip6*, Δ*SMDB11_0810-0809* (Δ*0810*Δ*0809*), and Δ*ssp6*Δ*sip6* Δ*SMDB11_0810-0809* (Δ*ssp6*Δ*sip6* Δ*0810*Δ*0809*) when co-cultured with wild type (WT) or mutant (Δ*tssE*, Δ*ssp6*, Δ*SMDB11_0810* and Δ*ssp6*Δ*SMDB11_0810*) attacker strains. Individual data points are overlaid with the mean +/- SEM (n = 5 biological replicates). (c) Membrane potential and permeability of cells of *E. coli* MG1655 carrying empty vector control (VC, pBAD18-Kn) or plasmids directing the expression of Ssp6 fused with an N-terminal OmpA signal peptide (sp-Ssp6), alone or with Sip6, or of SMDB11_0810 fused with a signal peptide (sp-0810), alone or with SMDB11_0809, was determined. Cells were grown in LB containing 0.2% L-arabinose, stained with DiBAC4(3) and propidium iodide and analysed by flow cytometry as in Figure 3. Bars show mean +/- SEM, with individual data points superimposed (n = 3 independent experiments). (d) Measurement of NPN uptake by *E. coli* expressing sp-Ssp6 alone or with Sip6, or sp-0810 alone or with SMDB11_0809. NPN accumulation is expressed as arbitrary fluorescence units (AU) and bars show mean +/- SEM, with individual data points superimposed (n=3 independent experiments).

**Supplementary Figure 8**

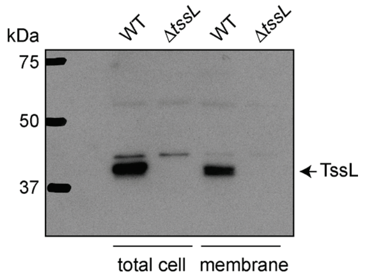


**Supplementary Figure 8. Validation of custom anti-TssL antibody.** Total cellular protein and total membrane protein samples were prepared from the wild type (WT) and a Δ*tssL* mutant of *S. marcescens* Db10, separated by SDS-PAGE and subjected to immunoblotting with a rabbit polyclonal antibody raised against the purified periplasmic domain of TssL (SMDB11_2254).

**Supplementary Table 1. Strains and Plasmids used in this study.**

| Name | Description/ genotype | Source / Reference |
|---|---|---|
| **Strains** | | |
| *Serratia marcescens* | | |
| Db10 | Wild type | [1] |
| SJC11 | Db10 Δ*tssE* (*SMDB11_2271*) | [2] |
| SJC3 | Db10 Δ*tssH* (*SMDB11_2274*) | [2] |
| KT74 | Db10 Δ*ssp6* (*SMBD11_4673)* | This study |
| KT75 | Db10 Δ*ssp6*Δ*sip6* (*SMBD11_4673, SMBD11_4672A)* | This study |
| KT77 | Db10 Δ*ssp6*Δ*sip6*, Sm-resistant derivative | This study |
| KT121 | Db10 Ssp6-HA (encodes Ssp6 [*SMDB11_4673*] with a C-terminal HA tag at the native chromosomal location) | This study |
| KT123 | Db10 Ssp6-HA, Δ*tssE* | This study |
| KT101 | Db10 Sip6-FLAG (encodes Sip6 [*SMDB11_4672A*] with a C-terminal 3xFLAG tag at the native chromosomal location) | This study |
| GM021 | Db10 Ssp6-HA, Sip6-FLAG | This study |
| AO07 | Db10 Δ*lacZ*::$P_{T5}$-*mCherry-kn*$^R$ (*lacZ*, *SMDB11_2462*). Encodes cytoplasmic mCherry, IPTG-inducible and expressed constitutively at a low level. | This study |
| AO08 | Db10 Δ*tssE*, Δ*lacZ*::$P_{T5}$-*mCherry-kn*$^R$ | This study |
| GM24 | Db10 Δ*ssp6*, Δ*lacZ*::PT5-*mCherry-kn*$^R$ | This study |
| SAN195 | Db10 Δ*lacZ*::$P_{T5}$-*gfpmut2-kn*$^R$. Encodes cytoplasmic GFP, IPTG-inducible and expressed constitutively at a low level. | [3] |
| GM04 | Db10 Δ*ssp6*Δ*sip6*, Δ*lacZ*::$P_{T5}$-*gfpmut2-kn*$^R$. | This study |
| GM143 | Db10 Δ*0810* (*SMDB11_0810)* | This study |
| GM141 | Db10 Δ*0810*Δ*0809* (*SMDB11_0810, SMDB11_0809*) | This study |
| GM144 | Db10 Δ*ssp6*, Δ*0810* | This study |
| GM142 | Db10 Δ*ssp6*Δ*sip6*, Δ*0810*Δ*0809* | This study |
| GM145 | Db10 Δ*0810*Δ*0809*, Sm-resistant derivative | This study |
| GM146 | Db10 Δ*ssp6*Δ*sip6,* Δ*0810*Δ*0809*, Sm-resistant derivative | This study |
| *Escherichia coli* | | |
| MG1655 | Wild type (model K-12 strain) | [4] |
| C43(DE3) | Protein overexpression strain; chromosomal λDE3 encodes IPTG-inducible T7 RNA polymerase F$^-$ *ompT hsdS* ($r^{B-}m^{B-}$) *gal dcm* λ(DE3) | Lucigen |
| CC118λ*pir* | Cloning host and donor strain for pKNG101-derived allelic exchange plasmids (λ*pir*) | [5] |
| HH26 pNJ5000 | Mobilizing strain for conjugal transfer | [6] |
| **Plasmids** | | |
| pSUPROM | Vector for constitutive expression of cloned genes under the control of the *E. coli tat* promoter (Kn$^R$) | [7] |
| pBAD18-Kn | Arabinose-inducible expression vector; gene of interest is cloned downstream of the $P_{ara}$ promoter (Kn$^R$) | [8] |
| pKNG101 | Suicide vector for allelic exchange (Sm$^R$, *sacBR, mobRK2, oriR6K*) | [9] |
| pNIFTY-MBP | Expression vector for T7 polymerase-dependent expression of recombinant proteins fused with N-terminal MBP-His$_6$ (Ap$^R$) | [10] |
| pSC1273 | Coding sequence for Sip6-HA (SMDB11_4672A) in pSUPROM | This study |

| | | |
|---|---|---|
| pSC1235 | Coding sequence for Ssp6 (SMDB11_4673) in pBAD18-Kn | [11] |
| pSC1236 | Coding sequence for $OmpA_{SP}$-Ssp6 fusion protein in pBAD18-Kn | [11] |
| pSC1256 | Coding sequences for Ssp6 + Sip6 in pBAD18-Kn | This Study |
| pSC1271 | Coding sequence for $OmpA_{SP}$-Ssp6 + Sip6 in pBAD18-Kn | This Study |
| pSC1587 | Coding sequence for $OmpA_{SP}$-MBP-Ssp6 + Sip6 in pBAD18-Kn | This Study |
| pSC1597 | Coding sequence for $OmpA_{SP}$-MBP-Ssp6 in pBAD18-Kn | This Study |
| pSC2549 | Coding sequence for $OmpA_{SP}$-SMBD11_0810 in pBAD18-Kn | This Study |
| pSC2561 | Coding sequence for $OmpA_{SP}$-SMBD11_0810 + SMDB11_0809 in pBAD18-Kn | This Study |
| pSC1270 | Coding sequence for Ssp6 in pNIFTY-MBP | This study |
| pSC1264 | pKNG101-derived allelic exchange plasmid for the generation of chromosomal in-frame Δ*ssp6* deletion | This study |
| pSC1266 | pKNG101-derived allelic exchange plasmid for the generation of chromosomal in-frame Δ*ssp6*Δ*sip6* deletion | This study |
| pSC1311 | pKNG101-derived allelic exchange plasmid for the generation of Ssp6-HA allele at the normal chromosomal location | This study |
| pSC1290 | pKNG101-derived allelic exchange plasmid for the generation of Sip6-3xFLAG allele at the normal chromosomal location | This study |
| pSC1509 | pKNG101-derived allelic exchange plasmid for the generation of Sip6-3xFLAG allele at the normal chromosomal location of strains harbouring Ssp6-HA | This study |
| pSC2553 | pKNG101-derived allelic exchange plasmid for the generation of chromosomal in-frame Δ*SMDB11_0810* deletion | This study |
| pSC2554 | pKNG101-derived allelic exchange plasmid for the generation of chromosomal in-frame Δ*SMDB11_0810* Δ*SMD11_0809* deletion | This study |
| pSC1706 | pKNG101-derived allelic exchange plasmid for the generation of chromosomal Δ*lacZ*::$P_{T5}$-*mCherry*-$kn^R$ | This study |

#### Supplementary Table 2. Oligonucleotide primers and additional details for plasmid construction

| Plasmid | Sequence of relevant primers (5'-3')[a] | Description |
|---|---|---|
| pSC1264 | TATATCTAGAGGGATCAGTTCGATGTGCG | Forward primer to clone upstream region of SMDB11_4673 in pKNG101 (*Xba*I) |
| | TATAAAGCTTACCTTTTGCCATGACTCAATTCC | Reverse primer to clone upstream region of SMDB11_4673 in pKNG101 (*Hin*dIII) |
| | TATAAAGCTTTTTGAAAAGGTGTCGAAATGAAGG | Forward primer to clone downstream region of SMDB11_4673 in pKNG101 (*Hin*dIII) |
| | TATAGGGCCCCGACGACATCAAGTACCTGTTCG | Reverse primer to clone downstream region of SMDB11_4673 in pKNG101 (*Apa*I) |
| pSC1266 | TATATCTAGAGGGATCAGTTCGATGTGCG | Forward primer to clone upstream region of SMDB11_4673 and SMDB11_4672A in pKNG101 (*Xba*I) |
| | TATAAAGCTTACCTTTTGCCATGACTCAATTCC | Reverse primer to clone upstream region of SMDB11_4673 and SMDB11_4672A in pKNG101 (*Hin*dIII) |
| | TATAAAGCTTGGCGTTTAGAGCAAGCGTAGAG | Forward primer to clone downstream region of SMDB11_4673 and SMDB11_4672A in pKNG101 (*Hin*dIII) |
| | TATAGGGCCCGCAGCGCAAGAAAGACGG | Reverse primer to clone downstream region of SMDB11_4673 and SMDB11_4672A in pKNG101 (*Apa*I) |
| pSC1290 | TATATCTAGACCAAGAGATGGCGAACGC | Forward primer to amplify SMDB11_4672A for generation of Sip6-3xFLAG fusion construct by overlap PCR (*Xba*I) |
| | CACCGTCATGGTCTTTGTAGTCAACGCCATCATGGAAC TTC | Reverse primer to amplify SMDB11_4672A for generation of Sip6-3xFLAG fusion construct by overlap PCR |
| | GAAGTTCCATGATGGCGTTGACTACAAAGACCATGAC GGTG | Forward primer to amplify 3x-FLAG tag (see below) for generation of Sip6-3xFLAG fusion construct by overlap PCR |
| | GAACGCTCTACGCTTGCTCTACTTGTCATCGTCATCCT TGTAATC | Reverse primer to amplify 3x-FLAG tag for generation of Sip6-3xFLAG fusion construct by overlap PCR |
| | GATTACAAGGATGACGATGACAAGTAGAGCAAGCGTA GAGCGTTC | Forward primer to amplify SMDB11_4672A downstream region for generation of Sip6-3xFLAG fusion construct by overlap PCR |
| | TATAGGGCCCGCAGCGCAAGAAAGACGG | Reverse primer to amplify SMDB11_4672A downstream region for generation of Sip6-3xFLAG fusion construct by overlap PCR (*Apa*I) |
| pSC1509 | TATATCTAGAGTCGATGTAGATAAAGTCAAAAGATG | Forward primer to amplify SMDB11_4672A for generation of Sip6-3xFLAG fusion construct by overlap PCR (*Xba*I), to incorporate 3xFLAG tag in strains harbouring Ssp6-HA |
| | Other primers as pSC1290 | |
| pSC2553 | TATATCTAGACACGCGCTTCGGCGGTATAAC | Forward primer to clone upstream region of SMDB11_0810 in pKNG101 (*Xba*I) |
| | TATAAAGCTTGGCATCCGCCATGTCGT | Reverse primer to clone upstream region of SMDB11_0810 in pKNG101 (*Hin*dIII) |
| | TATAAAGCTTTTTTGCAAAGGTTAACGTATGAAGAAC | Forward primer to clone downstream region of SMDB11_0810 in pKNG101 (*Hin*dIII) |
| | TATAGTCGACCAGCGATCATGGCTATGC | Reverse primer to clone downstream region of SMDB11_0810 in pKNG101 (*Sal*I) |
| pSC2554 | TATATCTAGACACGCGCTTCGGCGGTATAAC | Forward primer to clone upstream region of SMDB11_0810 and SMDB11_0809 in pKNG101 (*Xba*I) |
| | TATAAAGCTTGGCATCCGCCATGTCGT | Reverse primer to clone upstream region of SMDB11_0810 and SMDB11_0809 in pKNG101 (*Hin*dIII) |
| | TATAAAGCTTGATTAAATAACATTGTGTTC | Forward primer to clone downstream region of SMDB11_0810 and SMDB11_0809 in pKNG101 (*Hin*dIII) |

| | | |
|---|---|---|
| | TATAGTCGACCAACGGCTATCAAAAGGTC | Reverse primer to clone downstream region of SMDB11_0810 and SMDB11_0809 in pKNG101 (*Apa*I) |
| pSC1270 | TATAGGCGCCATGGCAAAAGGTGCGAAGG | Forward primer to clone SMDB11_4673 in pNIFTY-MBP (*Kas*I) |
| | TATAAAGCTTCATTTCGACACCTTTTCAAAAAATAGC | Reverse primer to clone SMDB11_4673 in pNIFTY-MBP (*Hin*dIII) |
| pSC1273 | TATAGGATCCATGAAGGTTTTTTCAGTGCTCATATCAAG | Forward primer to clone SMDB11_4672A in pSUPROM (*Bam*HI) |
| | TATAGTCGACTTATGCATAATCAGGAACATCATAAGGATAAACGCCATCATGGAACTTCATG | Reverse primer to clone SMDB11_4672A in pSUPROM, incorporating a C-terminal HA tag (*SalI*) |
| pSC1256 | TATAGAATTCTCCAACGCCCCCTACG | Forward primer to clone SMDB11_4673, including RBS, and SMDB11_4672A in pBAD18-Kn (*Eco*RI) |
| | TATATCTAGAGTGAACTGAAAGCCTTCCGG | Reverse primer to clone SMDB11_4673 and SMDB11_4672A in pBAD18-Kn (*Xba*I) |
| pSC1271 | TATATCTAGAGCAAAAGGTGCGAAGGAAATC | Forward primer to clone SMDB11_4673 and SMDB11_4672A into pSC1236 for fusion of SMDB11_4673 with OmpA$_{sp}$ (*Xba*I) |
| | TATAGTCGACTGCTCTAAACGCCATCATGG | Reverse primer to clone SMDB11_4673 and SMDB11_4672A in pSC1236 (*Sal*I) |
| pSC1587 | TATAGTCGACGAAAAGGTGTCGAAATGAAGGTTTTTTCAGTGCTC | Forward primer to clone SMDB11_4672A into pSC1597 (*Sal*I) |
| | TATAGCATGCCTAAACGCCATCATGGAACTTC | Reverse primer to clone SMDB11_4672A into pSC1597 (*SphII*) |
| pSC1597 | TATATGTAGAATGAAAATCGAAGAAGGTAAACTG | Forward primer to amplify SMDB11_4673 incorporating a N-terminal MBP tag from pSC1270, and clone into pSC1236 (*Xba*I) |
| | TATAGTCGACTCATTTCGACACCTTTTCAAAAAATAGC | Reverse primer to amplify SMDB11_4673 incorporating a N-terminal MBP tag from pSC1270, and clone into pSC1236 (*Sal*I) |
| pSC2549 | TATATCTAGAGCGGATGCCGTAAACATTGACAGC | Forward primer to clone OmpA$_{sp}$SMDB11_0810 in pBAD18-Kn (*Xba*I) |
| | TATAGTCGACTTATGCATAATCAGGAACATCATAAGGATAAACGCCATCATGGAACTTCATG | Reverse primer to clone OmpA$_{sp}$SMDB11_0810 in pBAD18-Kn (*Sal*I) |
| pSC2561 | TATAGTCGACAGAGGACGTTAAAATGAAGAACTTTATGCTG | Forward primer to clone SMDB11_0809 incorporating RBS from SMDB11_2264 in pSC2549 (*Sal*I) |
| | TATAGTCGACTCATACGTTAACCTTTGC | Reverse primer to clone SMDB11_0809 incorporating RBS from SMDB11_2264 in pSC2549 (*Pst*I) |

| **Plasmid** | **Relevant information** |
|---|---|
| pSC1290 & pSC1509 | Nucleotide sequence of 3xFLAG tag:<br>GACTACAAAGACCATGACGGTGATTATAAAGATCATGATATCGATTACAAGGATGACGATGACAAGTAG |
| pSC1706 | Synthetic insert produced by Invitrogen GeneArt (ThermoFisher) containing the mCherry sequence from pmCherry-N1 and used to replace P$_{T5}$-*gfpmut2* with P$_{T5}$-*mCherry* (*Bam*HI and *Sal*I) in pSAN72[3] |
| pSC1311 | Synthetic insert produced by Invitrogen GeneArt (ThermoFisher):<br>Contains sequence encoding the final 206 amino acids of SMDB11_4673 (not including the stop codon) immediately followed by an HA tag (YPYDVPDYA) then the final five amino acids and stop codon of SMDB11_4673 (repeated to preserve translation of the overlapping downstream gene) and then the following 596 nucleotides downstream of SMDB11_4673 (*Xba*I-*Apa*I) |

[a]Incorporated restriction sites for cloning into the respective vector are underlined.
